# The dual Ras Association (RA) Domains of Drosophila Canoe have differential roles in linking cell junctions to the cytoskeleton during morphogenesis

**DOI:** 10.1101/2024.07.29.605598

**Authors:** Emily D. McParland, Noah J. Gurley, Leah R. Wolfsberg, T. Amber Butcher, Abhi Bhattarai, Corbin C. Jensen, Ruth I. Johnson, Kevin C. Slep, Mark Peifer

## Abstract

During embryonic development and adult homeostasis epithelial cells must change shape and move without disrupting the tissue’s dynamic architecture. This requires robust linkage of cell-cell adherens junctions to the force-generating actomyosin cytoskeleton. Drosophila Canoe and mammalian Afadin play key roles in this linkage. One central task for the field is defining how upstream inputs from Ras-family GTPases regulate Canoe and Afadin. They are unusual in that they share two tandem Ras-association (RA) domains, which, when deleted, virtually eliminate Canoe function. Previous work in vitro suggested RA1 and RA2 differ in their ability to bind GTPases, but their individual functions in vivo remain unknown. Combining bioinformatic and biochemical approaches, we find that both RA1 and RA2 bind to active Rap1 with similar affinities, and that conserved N-terminal extensions play a role in binding. We created Drosophila *canoe* mutants to test RA1 and RA2 function in vivo. Despite their similar affinities for Rap1, RA1 and RA2 play strikingly different roles. Deleting RA1 virtually eliminates Canoe function in morphogenesis, while mutants lacking RA2 are viable and fertile but have defects in junctional reinforcement in embryos and during pupal eye development. These data significantly expand our understanding of how adherens junction:cytoskeletal linkage is regulated.

## Introduction

Small GTPases of the Ras superfamily regulate a remarkably diverse array of cellular processes from signaling by Receptors tyrosine kinases to protein trafficking to cytoskeletal regulation (reviewed in (Cherfils and Zeghouf, 2013). One key challenge for the field is uncovering the mechanisms by which these GTPases regulate their protein effectors. Members of the Ras family, including human K-, N-, and H-Ras as well as Rap1, bind many regulators and effectors. Many bind via ubiquitin-fold Ras association (RalGDS/AF6 Ras associating; RA) domains (Smith, 2023; Wohlgemuth et al., 2005). Defining the specificity of these interactions and the biological consequences of effector binding are current goals.

We focus on mechanisms regulating cell shape change and movement during morphogenesis and the network of proteins that mediate this by linking cell-cell adherens junctions (AJs) to actin and myosin (Perez-Vale and Peifer, 2020). Drosophila Canoe (Cno) and its mammalian homolog Afadin are junctional adapter proteins that are key components of this network. They regulate diverse events ranging from initial apical positioning of AJs (Choi et al., 2013) to the cell shape changes of gastrulation and convergent elongation (Sawyer et al., 2011; Sawyer et al., 2009) to the collective cell migration events driving dorsal closure and head involution (Boettner et al., 2003; Choi et al., 2011). Cno and Afadin are complex multidomain proteins, sharing five folded protein domains and a long C-terminal intrinsically disordered region (Fig. 1A; (Gurley et al., 2023). The most N-terminal of these folded domains are two RA domains, consistent with idea that these proteins act as effectors of Ras family GTPases.

**Fig. 1.**
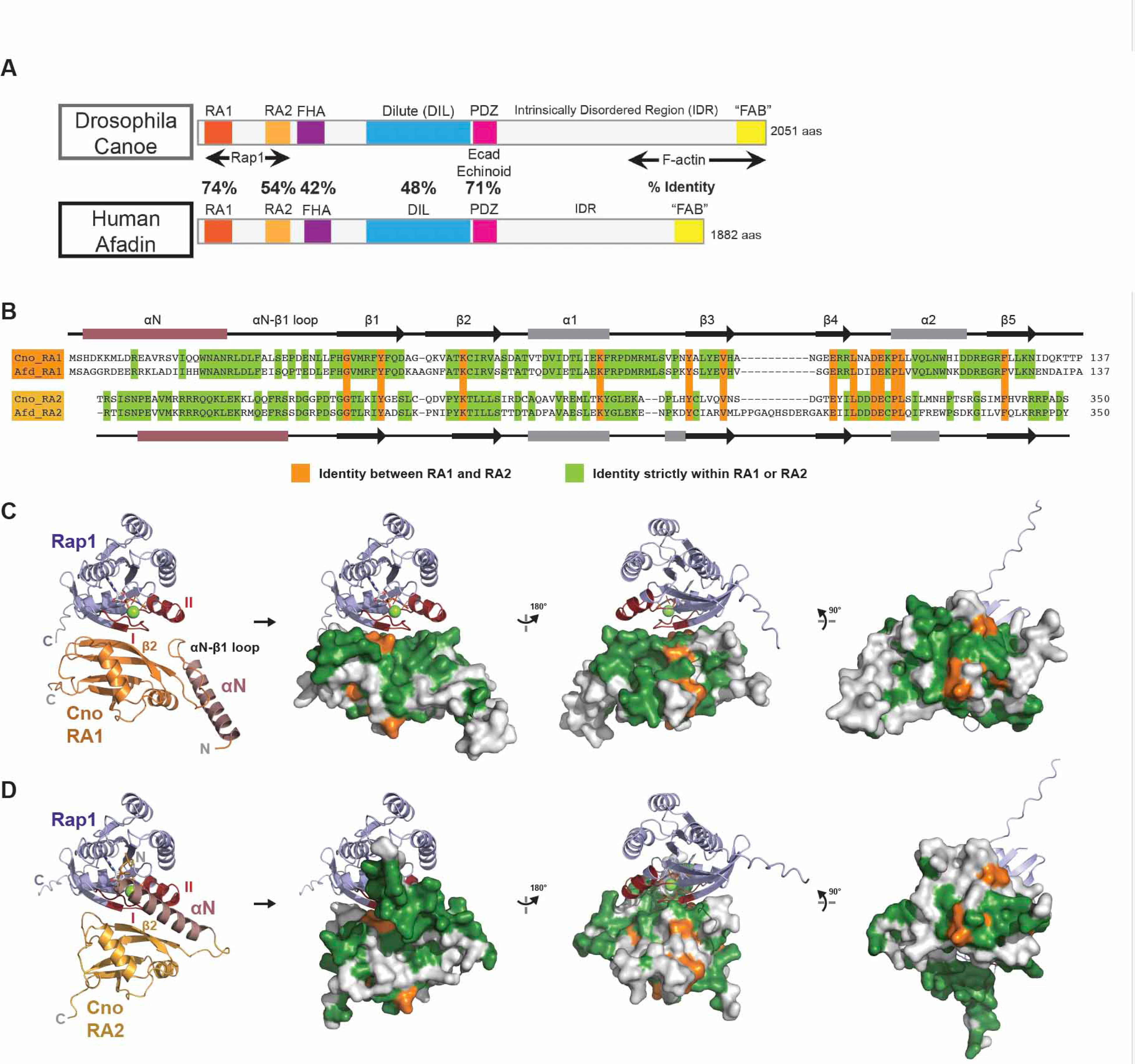
Cno RA domains are predicted to bind Rap1 using a conserved surface and display unique conserved surfaces outside the Rap1 binding site. (A) Diagrams comparing Drosophila Canoe and Human Afadin. (B) Sequence alignment of D.m. Canoe and mouse Afadin RA1 and RA2. Secondary structure is presented with αN colored in salmon. Residues identical strictly within either RA1 or RA2 are highlighted in green. Residues that are identical between RA1 and RA2 are highlighted in orange. (C,D) AlphaFold2 structure models of the D.m. Cno RA1:Rap1 complex (C) and the D.m. Cno RA2:Rap1 complex (D) shown in cartoon format at left, and with the RA domain in surface representation, with identity mapped as delineated in (B); right three panels, undergoing rotations as shown. The αN helix of each RA domain is colored salmon. Rap1 switch I and II regions are colored dark red. GMPPNP and Mg2+ are modeled based on alignments of the predictions with the Afadin RA1:Ras structure (PDB ID 6AMB, Fig. S1C; (Smith et al., 2017).

The Rap1 GTPase acts through multiple effectors to regulate cell-cell and cell-matrix adhesion during embryonic development and tissue homeostasis (Boettner and Van Aelst, 2009). Fly and human Rap1 share ∼55% identity to canonical Ras family members, and, in their activated state bind in vitro the RA domains of multiple effectors (Wohlgemuth et al., 2005). The list of Rap1 effectors includes a wide array of proteins, ranging from regulators of integrin-based adhesion, like Rap1-interacting adapter molecule (RIAM), to cytoskeletal regulators, like the Rac guanine nucleotide exchange factor (GEF) Tiam, to proteins that help link AJs to the actin cytoskeleton, like Cno and Afadin.

One way Afadin was originally identified was via biochemical searches for canonical Ras binding partners (Kuriyama et al., 1996; Van Aelst et al., 1994). Afadin can bind in vitro to multiple small GTPases including Rap1 (Boettner et al., 2000), M-Ras (Quilliam et al., 1999), Rit, and Rin (Shao et al., 1999). However, functional analysis in Drosophila revealed that Rap1 is a key regulator of Cno, in events ranging from apical-basal polarity establishment (Choi et al., 2013) to mesoderm invagination (Sawyer et al., 2009; Spahn et al., 2012) to dorsal closure (Boettner et al., 2003). Consistent with this, deleting both RA domains compromises all of Cno’s functions (Perez-Vale et al., 2021). Rap1 also acts through mammalian Afadin in multiple contexts from lumen formation in kidney tubules (Hiremath et al., 2023) to angiogenesis and barrier recovery after injury in the vascular endothelium (Birukova et al., 2013; Tawa et al., 2010), and deleting both RA domains reduces Afadin function in cultured MDCK cells (Choi et al., 2016). There is some evidence suggesting that other Ras family members may regulate Cno and Afadin. For example, R-ras regulates axon branching in cortical neurons via Afadin (Iwasawa et al., 2012) and Drosophila Ras may act through Cno in regulating cell fate or positioning in the developing eye (Gaengel and Mlodzik, 2003; Matsuo et al., 1997). However, the bulk of the evidence supports a key role for Rap1 in regulating Cno function and suggests that it acts, at least in part, via Cno’s RA domains (Bonello et al., 2018; Perez-Vale et al., 2021).

While most Ras family effectors and regulators have only a single RA domain, Cno/Afadin has two positioned in tandem (Fig. 1A). Given the essential role of the RA region in Cno function, it is now critical to explore the roles of each of the two RA domains in Rap1 binding and Cno function, to define whether they serve similar or different functions. The RA1 domain is very well conserved, with Drosophila and mammalian RA1 sharing 74% amino acid identity (Fig. 1A,B; (Gurley et al., 2023). Intriguingly, while most RA domains have a ββαββαβ ubiquitin fold (Mott and Owen, 2015), Afadin’s RA1 domain, like the RASSF5 RA domain, has an additional N-terminal alpha helix (Smith et al., 2017). Sequence conservation suggests this is shared by Cno RA1. RA2 is less well conserved, with 54% amino acid identity between Drosophila and mammalian proteins (Fig. 1A,B). Alphafold predictions (Jumper et al., 2021; Varadi et al., 2022) revealed that Afadin and Canoe RA2 also are likely to have an additional N-terminal alpha-helix (Fig. 1C). However, although both RA1 and RA2 are predicted to fold into similar structures, they are <u>very</u> divergent in sequence, with less than 30% sequence identity between Drosophila RA1 and RA2 (Fig. 1B).

Consistent with this, previous data suggested that their interactions with different Ras family members have diverged. Binding assays revealed that Afadin’s RA1 has a high affinity for Rap1 (Kd=0.24 µM), a lower affinity for H-Ras, MRas, or Rap2 (2-3 µM), and an even lower affinity for other Ras family members like R-Ras and TC21 (16 µM; (Linnemann et al., 1999; Wohlgemuth et al., 2005). In contrast, when these researchers measured binding of Afadin RA2 with small GTPases, they found a very low affinity for H-Ras (30 µM) and no detectable binding to Rap1 (Wohlgemuth et al., 2005). However, recent work suggests that these latter data may reflect an issue with the constructs used. The Ikura lab measured the affinity of Afadin RA1 with or without its N-terminal helix and found that deleting the N-terminal helix reduced affinity for Ras four-fold, from 4.1 µM to 17.8 µM (Smith et al., 2017). All measurements of Afadin RA2 interactions used a construct lacking the N-terminal helix while the parallel assays of RA1 used constructs that included the N-terminal helix (Wohlgemuth et al., 2005). It is possible RA1 and RA2 have specificities for different small GTPases, as pull down assays suggest Afadin RA2 may bind Rap2 (Goudreault et al., 2022). Analyses of Cno’s two RA domains have been less comprehensive. Yeast-two hybrid assays suggest that Rap1 interacts with both Cno RA1 and RA2—these data imply a stronger interaction with RA1 but affinities were not measured (Boettner et al., 2003). Despite these differences, the differential functions of the two RA domains have not been explored in either Cno or Afadin.

One other protein with two RA domains is Phospholipase C-epsilon (PLCε; (Dusaban and Brown, 2015), and in this protein the individual functions of each RA domain were explored in detail. PLCε integrates signaling from G-protein coupled receptors to downstream kinases via a catalytic role in generating lipid second messengers, and it also has a domain acting as a Rap1 GEF. It’s two RA domains are C-terminal. Like Afadin/Cno RA1, the RA2 domain of PLCε has clear small GTPase binding partners, including both the canonical Ras proteins and Rap1—canonical Ras proteins bind PLCε RA2 with ∼8-fold higher affinity than does Rap1 (Bunney et al., 2006). RA2 mediates both PLCε activation by Ras, Rap1 and their upstream activators and its recruitment to the plasma membrane (Bunney et al., 2006; Kelley et al., 2001; Song et al., 2002). In contrast, PLCε’s RA1 domain does not bind with high affinity to these same partners (Bunney et al., 2006). Instead, structural studies revealed that RA1 mediates intradomain interactions that promote the stability and basal activity of PLCε (Rugema et al., 2020).

Thus, multiple models of the roles of the RA1 and RA2 domains of Cno remain to be tested. They may be functionally redundant with similar affinities of small GTPases, they may have different affinities for Ras family members, or one may serve a more structural function. We thus tested the roles of Cno’s RA1 and RA2 domains, combining biochemical approaches to determine whether both can bind Rap1, and genetic and cell biological analyses of their functions in vivo.

## Results

### Conserved regions N-terminal to each RA domain are predicted to potentiate Rap1 binding

Drosophila Cno and mammalian Afadin share substantial sequence identity between their respective RA domains (74% RA1, 54% RA2, Fig. 1A,B, residues highlighted in green and orange). However, RA1 and RA2 are much less similar to one another, with only 14% amino acid identity over the canonical RA domain region (Fig 1B, residues highlighted in orange). As prior work reported a higher affinity between activated Rap1 and RA1, and weaker or no affinity for RA2, we used AlphaFold2 to generate models of the Cno RA1:Rap1 and Cno RA2:Rap1 complexes, to reveal potential structural attributes that may underlie differences in affinity, and to allow us to map potential functional regions shared by RA1 or RA2 domains. As the region N-terminal to each RA domain is conserved, and prior work revealed that a predicted helix N-terminal to RA1 increased the affinity for activated Ras (Smith et al., 2017), we included these regions in the sequences inputted to AlphaFold2. AlphaFold2 generated structural models with high confidence (Fig. S1A,B; see pLDDT plots), with Rap1 engaging both RA1 and RA2 using Switch regions I and II in their active GTP-bound state, and the C-terminal β-strand of Switch I engaging the β2 strand of each RA domain in an anti-parallel interaction. The predicted interaction aligns well with the solved crystal structure of mouse Afadin RA1 in complex with activated human Ras (RMSD = 0.70 Å over 147 Cα atoms, Fig. S1C, (Smith et al., 2017). The Cno RA1:Rap1 and Cno RA2:Rap1 complexes also align well with each other (RMSD = 1.08 Å over 204 Cα atoms, Fig. S1D).

Interestingly, each model included a predicted helix N-terminal to the RA domain that appear to support Rap1 binding in different ways. The αN helix preceding RA1 docks alongside the RA domain, stabilizing the conserved αN-β1 loop which in turn is modeled contacting the Rap1 Switch II region (Fig. 1C). While this N-terminal region was not fully present or ordered in the Afadin RA1 construct crystallized in complex with Ras, the model aligns with biochemical and biophysical data from the report that the αN helix potentiates Ras binding (Smith et al., 2017). The Cno RA2:Rap1 model also predicts a helix N-terminal to RA2 (Fig. 1D). The conformation of this αN helix is distinct from that of the RA1 αN helix. Consistent with this, there is no conserved sequence identity between the αN helix of RA2 and the αN-β1 loop of RA1 (Fig. 1B-D). In the Cno RA2:Rap1 model, the αN helix is not docked alongside the RA domain, but instead is modeled contacting the Rap1 Switch I region and may engage the GTP ribose (Fig. 1D). Mapping identity between or strictly within RA1 and RA2 on the structure models highlights distinct conservation at the Rap1 binding site, which could lead to differences in affinity (Fig. 1C,D, right three panels). RA domain-specific identity also maps to regions that expand beyond the Rap1 binding site, including the relative back face of the RA domain. The conserved regions of RA1 and RA2 are distinct from one another, which suggests that each RA domain may, by itself or when bound to a GTPase, engage distinct targets.

### Cno’s RA domains both bind activated Rap1 with similar affinities

While AlphaFold2 predicted robust models of these Cno RA:Rap1 complexes, we questioned how distinct conservation and unique N-terminal segments might affect Rap1 binding affinity, differences that might suggest unique roles in protein function. To test the relative binding activity of Cno RA1 and RA2 to activated Rap1, we employed two binding assays: Size exclusion chromatography with multiangle light scattering (SEC-MALS) and isothermal titration calorimetry (ITC). We purified the constitutively active G12V Rap1 mutant (which hydrolyzes GTP slowly) and further stabilized the active state by exchanging in the non-hydrolyzable nucleotide, GMPPNP. We first analyzed Rap1-GMPPNP and RA1 alone, and in complex using SEC-MALS. Each individual protein eluted as a single peak and produced an experimental mass near its formula mass (Fig 2A,A’). When we incubated stoichiometric amounts of Cno RA1 and Rap1-GMPPNP, a distinct peak eluted earlier than the individual proteins (15.0 ml point) with an experimental mass of 35.7 kDa, on par with a 1:1 complex formula mass of 36.1 kDa (Fig. 2A’’,A’’’). Some excess RA1 domain eluted at the 16.4 ml, likely due to errors in measuring stoichiometry (Fig. 2A’’). The robust elution shift of the complex, the profile of the peak, and the match between experimental and formula masses indicates that the interaction between RA1 and activated Rap1 is robust over the course of the gel filtration run (flow rate: 0.5 ml/min).

**Fig. 2.**
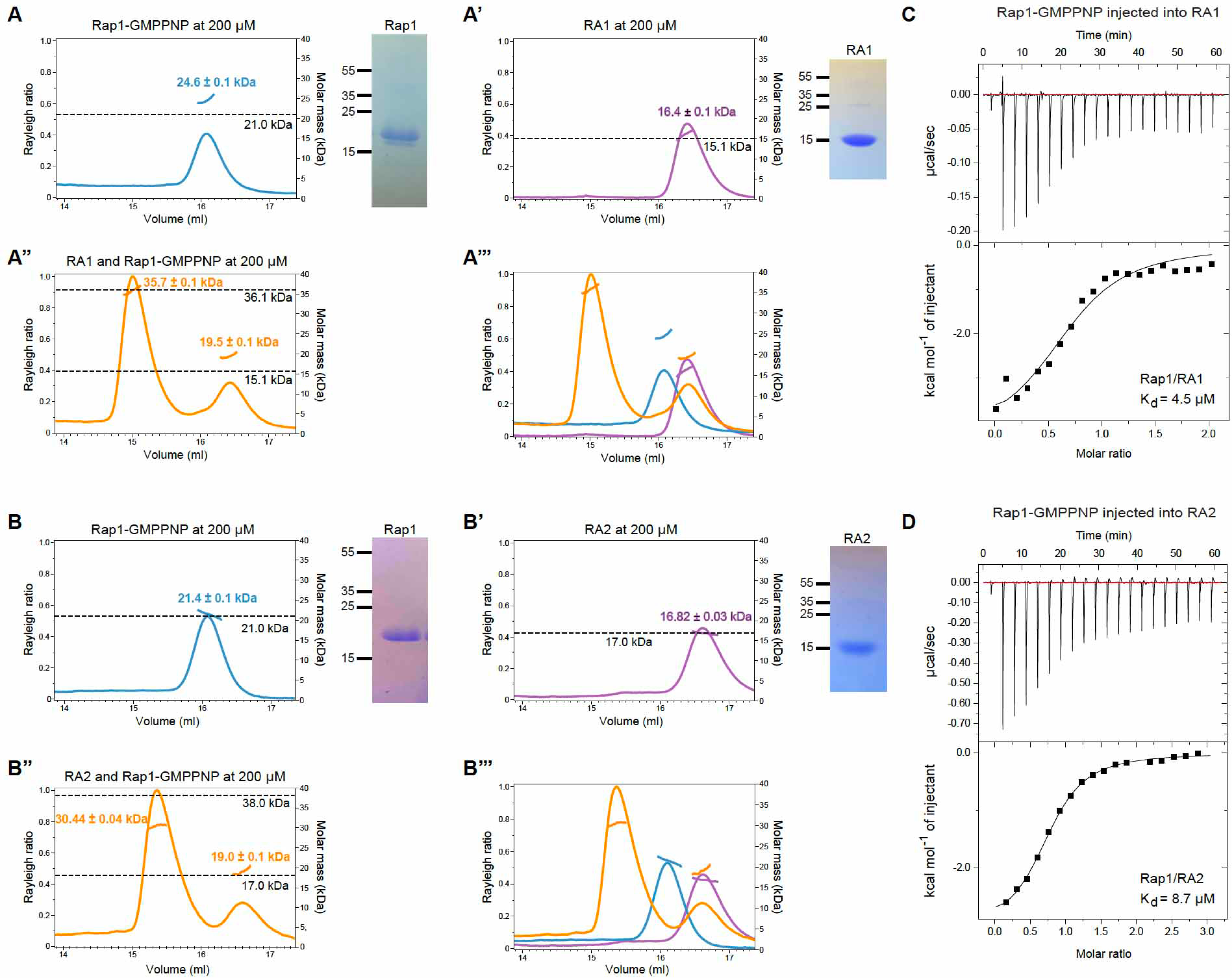
Activated Rap1 forms stable complexes with both RA1 and RA2. (A,B) SEC-MALS analysis of D.m. Rap1 and Cno RA domains alone and in complex. Formula molecular weights for GMPPNP-bound Rap1 G12V, RA1, and RA2 and 1:1 stoichiometric Rap1:RA domain complexes are indicated by dashed horizontal lines and were confirmed with Coomassie Blue-stained SDS PAGE. Experimental masses were determined by SEC-MALS. Each SEC-MALS experiment was performed in duplicate and a representative run is shown. (A) Rap1-GMPPNP-Mg has a formula mass of 21.0 kDa and an experimental mass of 24.6 ± 0.1 kDa. (A’) RA1 has a formula mass of 15.1 kDa (His tag removed) and an experimental mass of 16.4 ± 0.1 kDa. (A’’) The complex has a formula mass of 36.1 kDa and an experimental mass of 35.7 ± 0.1 kDa (left peak). Unbound RA1 domain (right peak) has an experimental mass of 19.5 ± 0.1 kDa. (A’’’) Super-positioning of the runs presented in A-A’’. (B) Rap1-GMPPNP-Mg has a formula mass of 21.0 kDa and an experimental mass of 21.4 ± 0.1 kDa. (B’) RA2 with a His tag has a formula mass of 17.0 kDa and an experimental mass of 16.82 ± 0.03 kDa. (B’’) The complex has a formula mass of 38.0 kDa and an experimental mass of 30.44 ± 0.04 kDa (left peak). Unbound RA2 domain (right peak) has an experimental mass of 19.0 ± 0.1 kDa. (B’’’) Super-positioning of the runs presented in B-B’’. Errors reported for SEC-MALS are SEM. (C,D) Isothermal titration calorimetry (ITC) analyses of the interactions between Rap1-GMPPNP and RA1 (C) and RA2 (D). Rap1-GMPPNP was injected into a well containing the respective RA domain over the course of one hour. Rap1-RA1 ITC experiments were performed in duplicate (biological): Expt. 1: Rap1/RA1 Kd = 4.5 µM, N = 0.717, ΔH = -4381 cal/mol; Expt. 2: Rap1/RA1 Kd = 5.9 µM, N = 1.00, ΔH = -4791 cal/mol. Rap1-RA2 experiments were performed in triplicate (a biological duplicate and a technical duplicate (technical duplicates: experiments 2 and 3): Expt. 1: Rap1/RA2 Kd = 8.7 µM, N = 0.755, ΔH = -3073 cal/mol. Expt. 2: Rap1/RA2 Kd = 8.8 µM, N = 0.73, ΔH = -2178 cal/mol. Expt. 3: Rap1/RA2 Kd = 7.9 µM, N = 0.63, ΔH = -2718 cal/mol.

We next performed a similar series of experiments using RA2 and Rap1-GMPPNP. Again, individual proteins eluted as single peaks and had experimental masses near to the formula mass (Fig 2B,B’). When we incubated stoichiometric amounts of Cno RA2 and Rap1-GMPPNP, an earlier-shifted peak again appeared (15.4 ml point) with an experimental mass of 30.4 kDa, larger than each individual component, but somewhat smaller than the complex formula mass of 38.0 kDa (Fig. 2B’’,B’’’). Some excess RA2 domain eluted at the 16.6 ml point, again likely due to errors in measuring stoichiometry (Fig. 2B’’). While the RA2-Rap1 complex produced a dramatic peak shift relative to the individual components, we note that the peak did not shift as much as the RA1-Rap1 complex, nor did the experimental mass fully match the formula mass of the complex. This may be due to some dissociation of the complex during the run, which would yield a mixed species and a lower apparent mass.

We next analyzed Rap1-GMPPNP binding affinity to each of Cno’s RA domains using ITC. Rap1-GMPPNP bound Cno RA1 with a Kd of 4.5 µM (Fig. 2C), and Cno RA2 with a Kd of 8.7 µM (Fig. 2D). The Rap1-RA1 affinity is on par with reported affinities between Afadin RA1 and Ras/Rap family small GTPases--e.g, Smith et al., (2017) reported a Kd=4.1 µM for Ras, while Wohlgemuth et al., (2005) reported a Kd=0.24 µM for Rap1 but 2-3µM for H-Ras, MRas, or Rap2. While the Rap1-RA2 affinity is about two-fold weaker in our hands than that of Rap1-RA1, it is still robust and greater than previously reported binding affinities using Afadin RA2 (Wohlgemuth et al., 2005). Our higher Rap1-RA2 affinity is likely due to the fact that we used a larger RA2 construct which includes the αN helix. Collectively, SEC-MALS and ITC binding studies demonstrate that Cno RA1 and RA2 can each bind activated Rap1 with micromolar affinity, and that RA1 has somewhat higher affinity for Rap1 than RA2. These findings align with predictions from the structural models in which common and distinct features of each RA domain are predicted to engage Rap1.

### Creating mutants individually deleting Cno’s two RA domains

Given the ability of each RA domain to bind Rap1, we returned to the question of whether they function redundantly or have unique functions. Deleting both RA domains severely reduced all the Cno functions we analyzed, altered Cno localization to nascent AJs, and eliminated Cno enrichment at cell-cell junctions under elevated tension (Perez-Vale et al., 2021). To define the functions of the individual RA domains, we generated mutants lacking one or the other RA domain—our deletions including the αN helix predicted to precede the respective RA domains (Fig. 3A, B). We have developed a system allowing us to replace the *cno* coding sequence at the endogenous locus with GFP-tagged proteins with specified mutational changes (Perez-Vale et al., 2021). We first designed mutants that cleanly deleted either the RA1 or RA2 domain, using the protein structures predicted by AlphaFold as a guide, creating *cnoΔRA1* and *cnoΔRA2*. In parallel, we generated mutants in which RA1 was cleanly replaced by a second copy of RA2 (*cnoRA2RA2*) and one in which RA2 was replaced by a second copy of RA1 (*cnoRA1RA1*; Fig. 3A, B). We verified each mutant by PCR and sequencing from the transgenic flies. We next examined protein accumulation, using both our polyclonal Cno antibody, which recognizes epitopes in the C-terminal IDR that are not altered in either mutant, or antibodies to the GFP tag. This verified existence of GFP-tagged proteins of the expected sizes, which accumulated in embryos at levels roughly similar to those of wildtype Cno (Fig. S2), as we observed with our previous mutant proteins (McParland et al., 2024; Perez-Vale et al., 2021). For CnoΔRA1 and CnoΔRA2 (Fig. S2A-F), which run at the same size as wildtype Cno (since the GFP tag and region deleted roughly match in size), we compared levels to those of our wildtype-GFP tagged Cno which accumulates at wildtype levels (Perez-Vale et al., 2021). CnoΔRA1 and CnoΔRA2 both accumulated at levels slightly lower than wildtype in early embryonic development and at levels slightly higher than wildtype later in embryonic development (Fig. S2E,F). CnoRA1RA1 and CnoRA2RA2 have a higher molecular weight than wildtype Cno, so we could directly compare levels of them to wildtype by comparing the two bands present in heterozygotes (Fig. S2G-L). Both accumulate at levels very close to wildtype (Fig. S2K,L). Finally, we outcrossed our mutants over multiple generations to a wildtype stock, removing any other potential mutations on the third chromosome.

**Fig 3.**
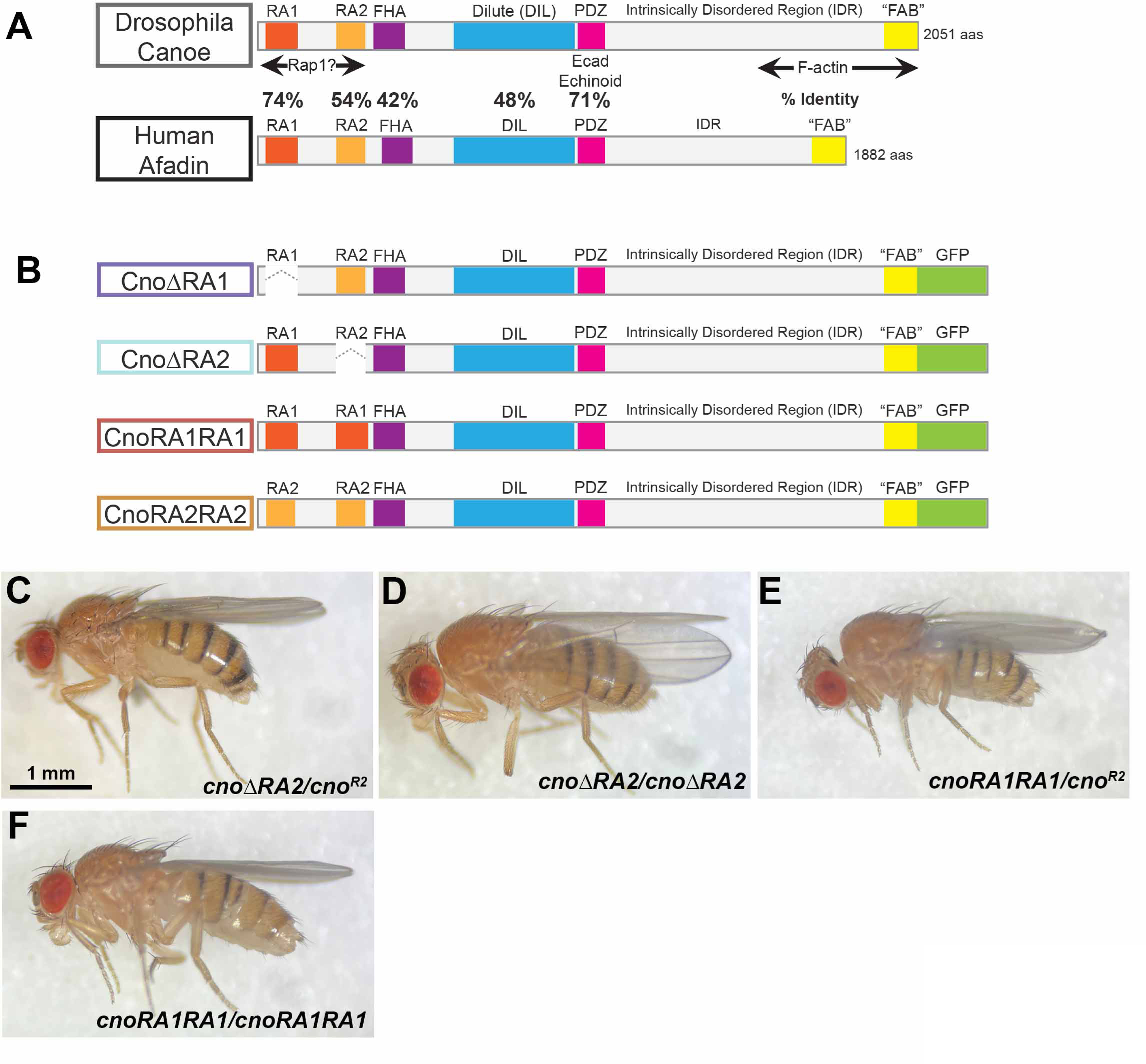
Generating mutants to test the function of Cno’s RA domains. (A) Diagrams comparing Drosophila Canoe and Human Afadin. (B) Diagrams of mutants examined. (C-F) Adult flies of the genotypes indicated. Both cno*ΔRA2* and cno*RA1RA1* are viable over the null allele, *cno^R2^,* and viable when homozygous.

### RA1 is essential for viability while RA2 is dispensable

For our initial tests of mutant protein function, we crossed each mutant to one of our canonical protein null alleles*, cno^R2^* (Sawyer et al., 2009), and assessed viability to adulthood. In this cross, *cnoΔRA* mutants, which lack both RA domains, are lethal before adulthood (Perez-Vale et al., 2021). Strikingly, while *cnoΔRA1* was lethal over *cno^R2^, cnoΔRA2/ cno^R2^* mutants were viable to adulthood at Mendelian ratios (Fig. 3C; 36% vs 33% predicted; n=736 adults). Consistent with this, we obtained homozygous *cnoΔRA2* adults and could create a homozygous mutant stock (Fig. 3D). *cnoΔRA2* maternal/zygotic mutant embryos had only 2% lethality (n=911 embryos), in the wildtype range. Thus, in this simple assessment of function, the RA1 domain is critical for viability, while the RA2 domain is dispensable.

We next asked whether adding a second copy of RA2 rescued the lethality resulting from loss of RA1, using the *cnoRA2RA2* mutant. *cnoRA2RA2* was also lethal over *cno^R2^.* In contrast, *cnoRA1RA1/ cno^R2^* mutants were viable to adulthood at Mendelian ratios (Fig. 3E; 36% vs 33% predicted; n=166), and we obtained homozygous *cnoRA1RA1* adults (Fig. 3F). They were fertile and *cnoRA1RA1* maternal/zygotic mutant embryos had 10% lethality (n=955), very slightly elevated from the wildtype range (generally 3-8% lethality). Thus, RA2 cannot replace RA1, while putting a second copy of RA1 in the place of RA2 has little or no effect on Cno function.

### RA1 is essential for embryonic morphogenesis

To fully assess CnoΔRA1 function, we needed to generate maternal/zygotic mutants. To do so, we used the FLP/FRT/ovoD approach (Chou and Perrimon, 1996) to generate females whose germlines were homozygous mutant for *cnoΔRA1* and crossed them to males heterozygous for *cnoΔRA1*. *cnoΔRA1* maternal/zygotic mutants were fully embryonic lethal, while most embryos receiving a wildtype gene were paternally rescued (53% embryonic lethality; n=895). As a first assessment of the function of CnoΔRA1 in embryonic morphogenesis, we assessed the cuticle secreted by the developing larva. Its features allow one to examine both integrity of the epidermal epithelium, and whether key morphogenetic movements like dorsal closure and head involution were completed. In wildtype (Fig. 4A), the epidermis is intact without holes, completion of head involution leads to development of an intact head skeleton (Fig. 4A arrow), and completion of dorsal closure means the cuticle is sealed dorsally. In *cno* zygotic null mutants, maternally contributed Cno is sufficient for many morphogenetic events and most embryos have defects in head involution (as in Fig. 4B,C). In contrast, embryos maternally and zygotically mutant for the null allele *cno^R10^* have fully penetrant defects in both dorsal closure and head involution (as in Fig. 4E; 94% of dead embryos; Fig. 4G), and many also have defects in the integrity of the ventral epidermis (as in Fig. 4F; 68%; Fig. 4G; Gurley et al., 2023). *cnoΔRA* maternal/zygotic mutants also have a strong phenotype, with defects that are similar, though slightly less severe, than those of the null mutants--dorsal closure and head involution fail, but defects in ventral epidermal integrity are less frequent (Perez-Vale et al., 2021). *cnoΔRA1* maternal/zygotic mutants also had severe disruptions of morphogenesis, with a very penetrant failure of dorsal closure and head involution (89% of the dead embryos); 24% also had additional holes in the ventral epidermis (Fig. 4G). This suggests loss of RA1 leads to a very strong loss of Cno function, with some potential residual function remaining.

**Fig. 4.**
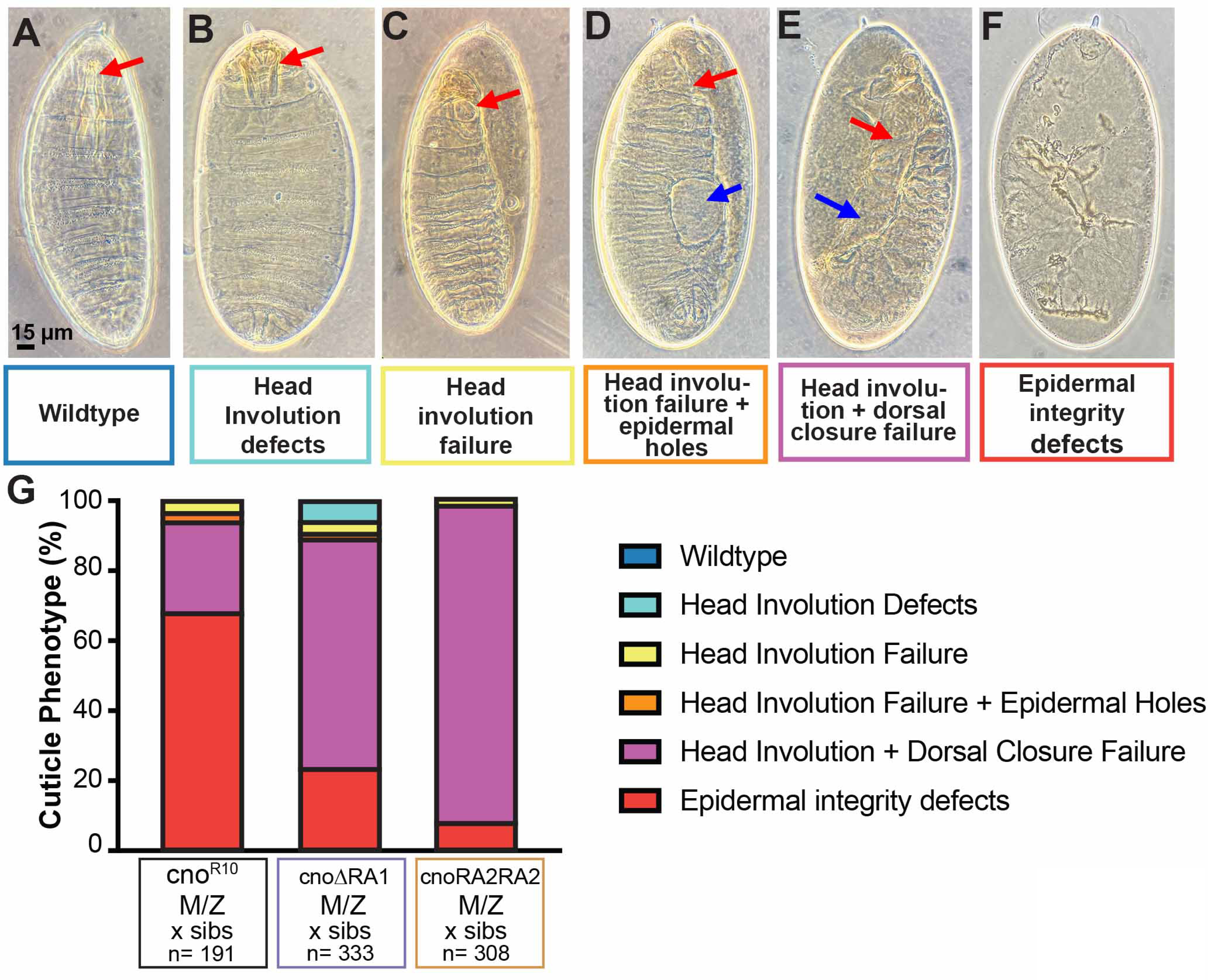
Deleting the RA1 domain severely reduces Cno function in embryonic morphogenesis. (A-F) Embryonic cuticles in the eggshell, illustrating possible defects differing in severity. (A) Wildtype with intact head skeleton (arrow). (B) Defects in head involution disrupt the head skeleton (arrow). (C). Complete failure of head involution, leading to an anterior hole (arrow). (D) Failure of head involution (red arrow) and holes in the dorsal or ventral epidermis (blue arrow). (E) Complete failure of head involution and dorsal closure, leaving the cuticle open anterior (red arrow) and dorsally (blue arrow). (F) More severe defects in epidermal integrity. (G) Stacked bar graph illustrating the range of defects in different mutants. In *cno* maternal/zygotic null mutants *(cno^R10^)*, most embryos combine complete failure of head involution and dorsal closure with additional defects in epidermal integrity. In contrast, most *cnoΔRA1* maternal/zygotic mutants are slightly less severe, with complete failure of head involution and dorsal closure but without additional epidermal holes. *cnoRA2RA2* maternal/zygotic mutants are similar but on average slightly less severe.

To test whether an additional copy of the RA2 domain compensates for loss of RA1, we next assessed CnoRA2RA2. We found that *cnoRA2RA2* maternal/zygotic mutants are fully embryonic lethal, while most embryos receiving a wildtype gene were paternally rescued (54% embryonic lethality; n=820). The cuticle phenotypes of the dead embryos revealed that CnoRA2RA2 also failed to rescue morphogenesis. Like *cnoΔRA1* mutants, they exhibited nearly fully penetrant failure of dorsal closure and head involution (95% of the dead embryos), and 9% also had holes in the ventral epidermis (Fig. 4G). Thus, replacing the RA1 domain with a second copy of the RA2 domain does not restore Cno function.

### Loss of RA1 disrupts Cno’s role in initial positioning of adherens junctions and dramatically reduces Cno recruitment to nascent junctions

Cno’s first role in morphogenesis occurs as cell-cell AJs are initially positioned apically during cellularization. As membranes invaginate around nuclei, the cadherin-catenin complex is enriched apically at nascent spot adherens junctions (SAJs Fig. 5A, B green arrowheads, C) along with Bazooka (Baz: fly Par3; (Harris and Peifer, 2004). There are also lower levels of the cadherin-catenin complex along the lateral membrane, and it is enriched at basal junctions (Hunter and Wieschaus, 2000) just behind the actin at the cellularization front (Fig. 5A, B red arrowheads). This is most apparent in maximum intensity projections (MIPs) of multiple cross sections (Fig. 5B). Cno localizes to the apico-lateral membrane where nascent AJs form (Fig. 5A,B; (Choi et al., 2013). Cno accumulates in all SAJs but is particularly enriched at tricellular junctions (TCJs; Fig. 5C, red vs green arrowheads; (Bonello et al., 2018). In the absence of Cno, both Armadillo (fly β-catenin) and Baz lose apical enrichment and localize all along the lateral membrane (Choi et al., 2013). Loss of Rap1 leads to total loss of Cno recruitment to the plasma membrane and mimics the effects of Cno loss on Arm and Baz localization (Choi et al., 2013). Surprisingly, deleting both RA domains led to an intermediate phenotype. Cno enrichment at TCJs was lost, Cno localization was not as tightly restricted to the region of the apical nascent junctions, and Arm and Baz enrichment at nascent apical junctions was reduced (Bonello et al., 2018; Perez-Vale et al., 2021).

**Fig. 5.**
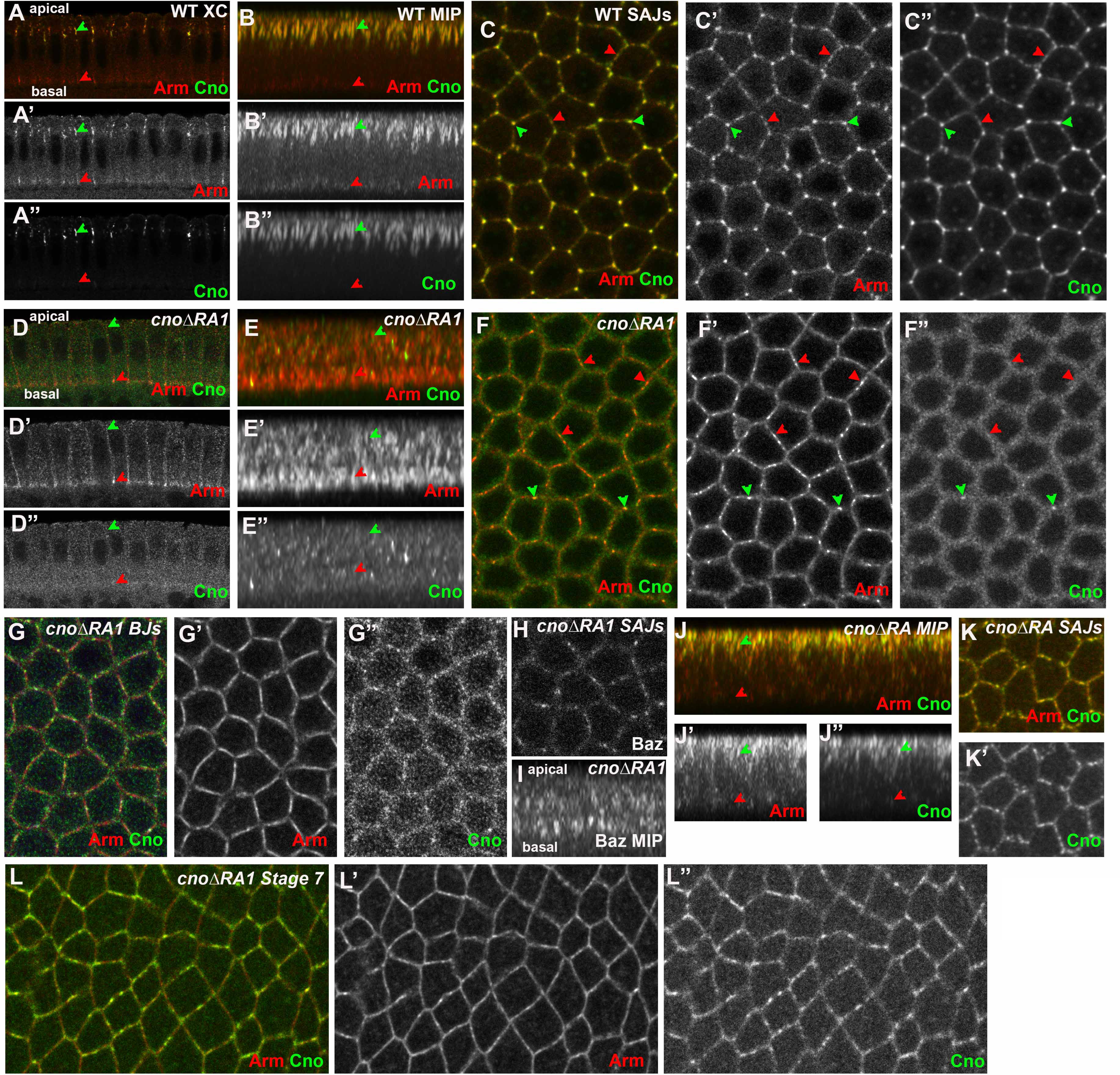
The RA1 domain is required for Cno localization and function as AJs are positioned during cellularization. (A-K) Late cellularization embryos, genotypes and antigens indicated. (A,B) Wildtype cross-section (A) and maximum intensity projection (MIP; B). Arm is enriched both in nascent apical junctions (green arrowheads) and in basal junctions (red arrowheads). Cno localizes to cables in apical junctions (green arrowheads) and is largely absent more basally (red arrowheads). (C) At the level of the apical spot AJs (SAJs) wildtype Cno is enriched at tricellular junctions (green arrowheads) relative to bicellular junctions (red arrowheads). (D,E) *cnoΔRA1* cross-section (D) and MIP (E). (F,G) Images at the level of the SAJs (F) and the basal junctions (G). Arm enrichment at apical junctions is reduced (D,E green arrowheads) while basal junction enrichment remains (D,E red arrowheads, G). CnoΔRA1 protein puncta are found all along the apical basal axis and membrane enrichment is reduced. No tricellular junction enrichment is apparent. (H,I) Baz at the level of the SAJs (H) and in MIP. The normal apical enrichment of Baz is lost. (J,K) In contrast, while CnoΔRA protein is less apically enriched (J), it remains localized to cell membranes. (L) As gastrulation starts, CnoΔRA1 protein returns to AJs.

We thus examined the localization and function of CnoΔRA1 during AJ establishment, using *cnoΔRA1* maternal/zygotic mutants. We were surprised to find that CnoΔRA1 was largely absent from the membrane (Fig. 5A” vs D”, B” vs E”, C” vs F”). Occasional Cno puncta accumulated along the lateral membrane, both at the level of the SAJs (Fig. 5F) and mislocalized to the level of the basal junctions (Fig. 5G). This is clearest in MIPs (Fig. 5E). This loss of membrane localization was consistently observed—in embryos stained in the same experiments 14/14 cellularizing embryos had little or no membrane localization of CnoΔRA1, while in 17/17 post-cellularization embryos CnoΔRA1 localized to cell junctions. This contrasts with CnoΔRA, lacking both RA domains, which remains enriched in membrane-proximal puncta (Fig. 5K; (Perez-Vale et al., 2021), though it loses its tight apical junctional restriction (Fig. 5J). *cnoΔRA1* mutants also fail to enrich Arm apically (Fig. 5D, E, green arrowheads), though some enrichment at basal junctions remains (Fig. 5D, E, red arrowheads). Baz also is no longer restricted to nascent apical junctions (Fig. 5I), although it continues to associate with the lateral membrane (Fig. 5H). Thus, CnoΔRA1 cannot support initial junctional polarization and has strongly reduced membrane localization—these data suggest CnoΔRA1 may be even less functional than CnoΔRA mutants, in which Arm localization to nascent junctions is reduced but not eliminated (Perez-Vale et al., 2021). Intriguingly, as gastrulation began during stage 6, CnoΔRA1 returned to apical junctions (Fig. 5L)—we observed a similar return of wildtype Cno to junctional localization at stage 6 in *Rap1* mutants (Bonello et al., 2018).

### Loss of RA1 destabilizes cell-cell junctions under elevated tension and alters junctional protein planar polarity

The morphogenetic movements of gastrulation, germband extension, and dorsal closure all require Cno function to stabilize AJs under elevated force as cells change shape. In its absence, AJs under elevated tension, like those at aligned anterior-posterior (AP) borders or at multicellular junctions, separate, and morphogenetic movements fail to go to completion. Since *cnoΔRA* is defective in all these Cno functions (Perez-Vale et al., 2021), we thus assessed morphogenetic movements, AJ stability and junctional protein planar polarity in *cnoΔRA1* maternal/zygotic mutants.

The first event of gastrulation is invagination of the mesoderm. In wildtype embryos the ectoderm then seals at the midline, leaving no gap (Fig. 6A). *cno* null mutants exhibit fully penetrant defects in this process, with mesodermal cells remaining on the surface and the furrow not fully closing (Sawyer et al., 2009). When we examined *cnoΔRA1* mutants (stages 7-9), 24/29 embryos had defects in mesoderm invagination, ranging from mild defects (Fig. 6B; 10/ 29 embryos) to more wide-open ventral furrows (Fig. 6C; 15/29). This was similar to the frequency of defects in *cnoΔRA* mutants (Perez-Vale et al., 2021).

**Fig. 6.**
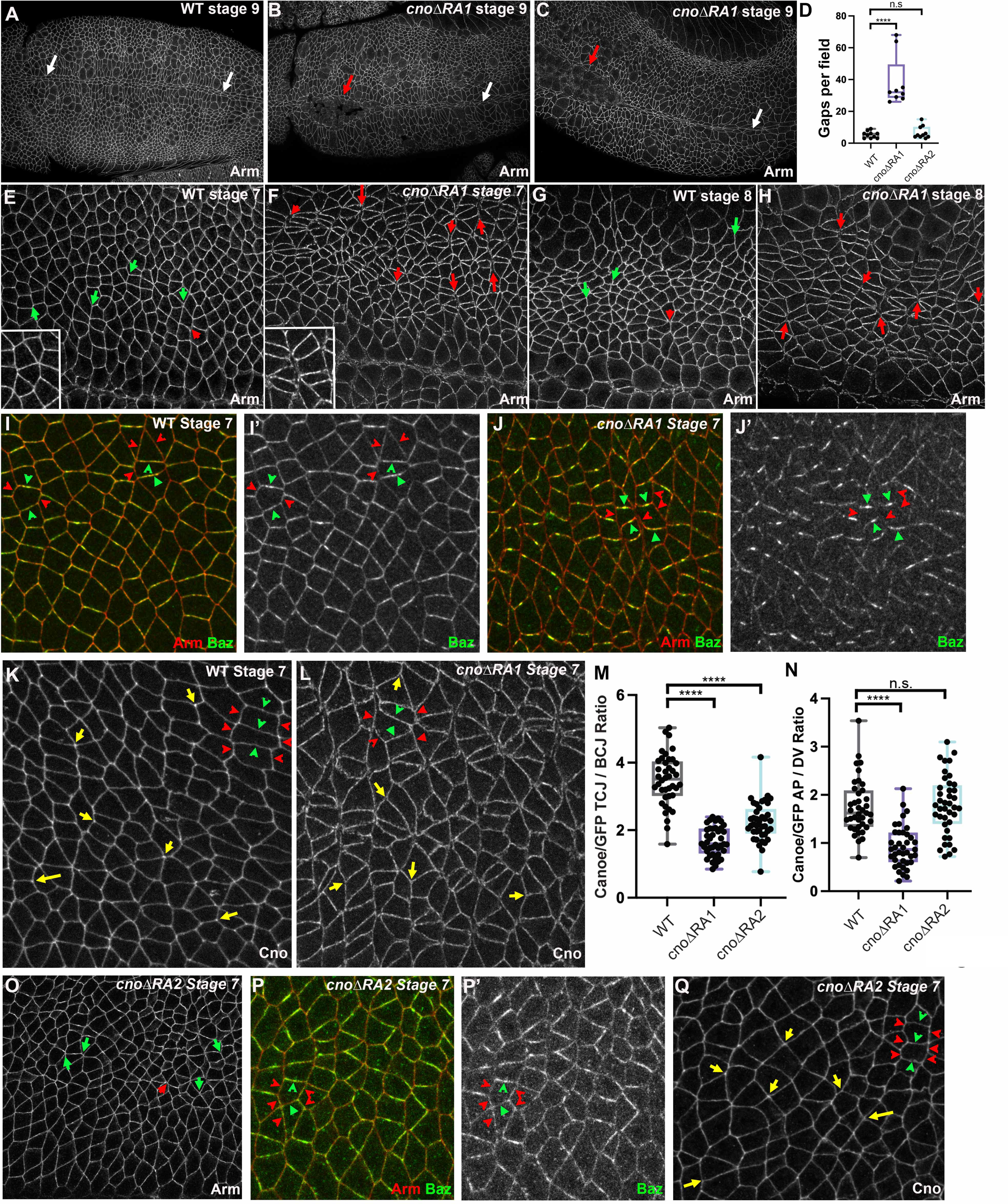
Cno’s RA1 domain is important for reinforcing AJs under tension and for Cno enrichment at these junctions. Embryos, anterior left and dorsal up, stage, genotype and antigens indicated. (A) In wildtype the ventral furrow is completely closed (arrows). (B,C) In many *cnoΔRA1* mutants, regions of the furrow have not closed (red arrows). (D) Quantification of gaps per field of view. (E,G) In wildtype at stages 7 and 8, very few gaps are seen in AJs (green vs red arrows). (F,H) In contrast, *cnoΔRA1* mutants have numerous gaps at aligned AP borders and rosette centers (red arrows). (I) In wildtype, Baz is subtly enriched at DV vs AP borders (green vs red arrowheads) but is found all around each cell. (J) In *cnoΔRA1* mutants Baz is substantially reduced at AP borders (red arrowheads) and often confined to the center of DV borders (green arrowheads). (K) In wildtype Cno is enriched at many TCJs (yellow arrows) and also is subtly enriched at AP vs DV borders (red vs green arrowheads). (L) In *cnoΔRA1* mutants both TCJ (yellow arrows) and AP border enrichment (red vs green arrowheads) are reduced. (M,N) Quantification of TCJ and AP border enrichment. (O) In *cnoΔRA2* mutants few gaps are seen in AJs (green vs red arrows). (P) Baz localization appears relatively normal in *cnoΔRA2* mutants. (Q) In *cnoΔRA2* mutants, some TCJ enrichment remains (yellow arrows) and enrichment at AP borders remains unchanged (red vs green arrowheads).

Germband extension is driven in part by reciprocal planar polarization of junctional and cytoskeletal proteins, with actin and myosin enriched on AP borders and AJ proteins and Baz enriched on dorsal-ventral (DV) borders. Junctional myosin contraction then shrinks AP borders and rearranges cells. We first examined whether the RA1 domain is essential to reinforce AJs under tension. As germband extension starts at stage 7 in wildtype, Arm localizes all around cells, with few apical gaps in AJs (Fig. 6E, green vs red arrows). In contrast, in *cnoΔRA1* mutants apical gaps appeared at many aligned AP borders and multicellular junctions (Fig. 6F, red arrows; Fig. E vs F, insets), similar to those we observed in *cnoΔRA* mutants (Perez-Vale et al., 2021). Similar gaps were seen at stage 8 (Fig. 6G vs H, red arrows), as cells in mitotic domain 11 rounded up to divide. Quantification verified a substantial increase in gap number in *cnoΔRA1* mutants (Fig. 6D; One-way ANOVA), with the frequency of gaps similar to that in *cnoΔRA* mutants (Perez-Vale et al., 2021). Cno is also important for restraining the planar polarization of Baz. In wildtype, Baz accumulates all around the cell, with ∼2-fold enrichment on DV vs AP borders (Fig. 6I, green vs red arrowheads). In Cno’s absence, Baz is reduced on AP borders and particularly enriched in the center of DV borders, elevating its planar polarization (Sawyer et al., 2011). *cnoΔRA1* mutants also had strong alterations in Baz localization, with very strong reduction of Baz on many AP borders (Fig. 6J, red arrowheads) and Baz often restricted to the central region of DV borders (Fig. 6J, green arrowheads). Again, this was similar to what was seen in both *cno* null mutants (Sawyer et al., 2011) and *cnoΔRA* mutants (Perez-Vale et al., 2021). Thus, the RA1 domain is critical for Cno reinforcement of AJs under elevated tension and for the restraint of Baz planar polarity—in these ways its mutant phenotypes closely resemble those of mutants lacking both RA domains (Perez-Vale et al., 2021).

### The RA1 domain is important for Cno enrichment at AJs under elevated tension

Cno protein is normally enriched at multicellular junctions (Fig. 6K, yellow arrows; quantified in Fig. 6M), and is also subtly enriched at AP borders (Fig. 6K, red vs green arrowheads; quantified in Fig. 6N), the same places where junctions apically separate in *cno* mutants (Perez-Vale et al., 2021). We next asked whether the RA1 domain is required for this enrichment. Inspection suggested reduced enrichment at multicellular junctions (Fig. 6L, yellow arrows) and loss of enrichment at AP relative to DV borders (Fig. 6L, red vs green arrowheads) in *cnoΔRA1* mutants. We quantified the ratio of Cno at tricellular to bicellular junctions, revealing that deleting the RA1 domain strongly reduces but does not eliminate tricellular junction enrichment (Fig. 6M). Quantification also verified loss of Cno enrichment at AP borders in *cnoΔRA1* mutants (Fig. 6N). Intriguingly, these phenotypes were similar to but not as strong as those of *cnoΔRA* mutants, which completely lose Cno enrichment at multicellular junctions and in which enrichment of Cno is reversed relative to wildtype, with elevated enrichment at DV borders (Perez-Vale et al., 2021).

### The RA2 domain is not essential for AJ stabilization

In parallel, we examined the role of the RA2 domain in stabilizing AJs under elevated tension, restraining planar polarization of Baz, and Cno localization to AJs under tension. When we examined *cnoΔRA2* mutants, we saw no significant increase in apical gaps in AJs relative to wildtype (Fig. 6O, quantified in 6D). Baz continued to localize to both AP and DV borders, without the extreme planar polarity seen in *cnoΔRA1* mutants (Fig. 6P’ vs J’). While CnoΔRA2 protein continued to be enriched at multicellular junctions (Fig. 6Q, yellow arrows), quantification revealed that average enrichment was reduced relative to wildtype, though not to the extent seen in *cnoΔRA1* mutants (Fig. 6M). Cno enrichment at aligned AP borders in *cnoΔRA2* mutants was similar to wildtype (Fig. 6N). Thus, consistent with its viability and fertility, the RA2 domain is not essential for most embryonic functions of Cno, though it may enhance enrichment at some AJs under elevated tension.

### The RA1 domain is essential for late morphogenesis and to maintain epidermal integrity

Cno is also essential for the later events of morphogenesis, including dorsal closure and head involution (Boettner et al., 2003; Jürgens et al., 1984). In *cno* maternal/zygotic null mutants both events fail, and embryos often exhibit additional issues with epidermal integrity. In these mutants, cells in the ventral epidermis have difficulty returning to columnar architecture after rounding up to divide, and this ultimately leads to holes in the ventral epidermis, which can be observed in the cuticle. We thus examined whether the RA1 domain was important for these Cno functions.

During stages 9 and 10 in wildtype, cells in the ectoderm can be divided into three groups. Dorsal ectodermal cells completed cell division in stage 8 and have since resumed columnar architecture. Cells in the lateral and ventral ectoderm divide during stages 9 and 10 and thus need to remodel junctions as they round up for mitosis and then resume columnar architecture. Simultaneously, a subset of these cells (∼30%) apically constrict and move inward to become neural stem cells. These twin challenges mean the ventral epidermis is more sensitive to disruptions that reduce adhesion or linkage of junctions to the cytoskeleton.

By stage 10 in wildtype, most cells have completed division, though a subset near the ventral midline remain rounded up in mitosis (Fig. 7A, white arrows). Cells rapidly resume columnar architecture after division. In contrast, in *cnoΔRA1* mutants, cells near the ventral midline fail to return to a columnar architecture (Fig. 7B, red arrows). By stage 11 in wildtype (Fig. 7C), cell division is largely complete, and cell shape changes are driving head segmentation and tracheal pit invagination. In *cnoΔRA1* mutants, groups of ventral cells that failed to return to a columnar architecture remain—the severity of this phenotype varies from moderate (Fig. 7D) to more severe (Fig. 7E). We also examined how AJ protein localization is altered in regions where epithelial architecture is altered. At stages 10 and 11 in wildtype, both Arm (Fig. 7I’) and Cno (Fig. 7J) are strongly enriched at AJs, with some modest reduction in junctional accumulation in cells rounded up to divide. Baz also localizes to AJs all around cells (Fig. 7I”). In *cnoΔRA1* mutants, epithelial cells that retain columnar architecture retain junctional Arm and Cno (Fig. 7K, yellow arrows), whereas cells that failed to resume columnar architecture had strongly reduced levels at AJs (Fig. 7K, cyan arrow). Baz localization to AJs was reduced in most columnar cells (Fig. 7K yellow arrows), and AJs appeared fragmented where columnar and non-columnar cells met, perhaps places where junctional remodeling is maximized. These junctional fragments accumulated all three AJ proteins (Fig. 7K, red arrows). These are all defects we have previously observed in our strongest *cno* mutants, including *cnoΔRA* (Manning et al., 2019; Perez-Vale et al., 2021).

**Fig. 7.**
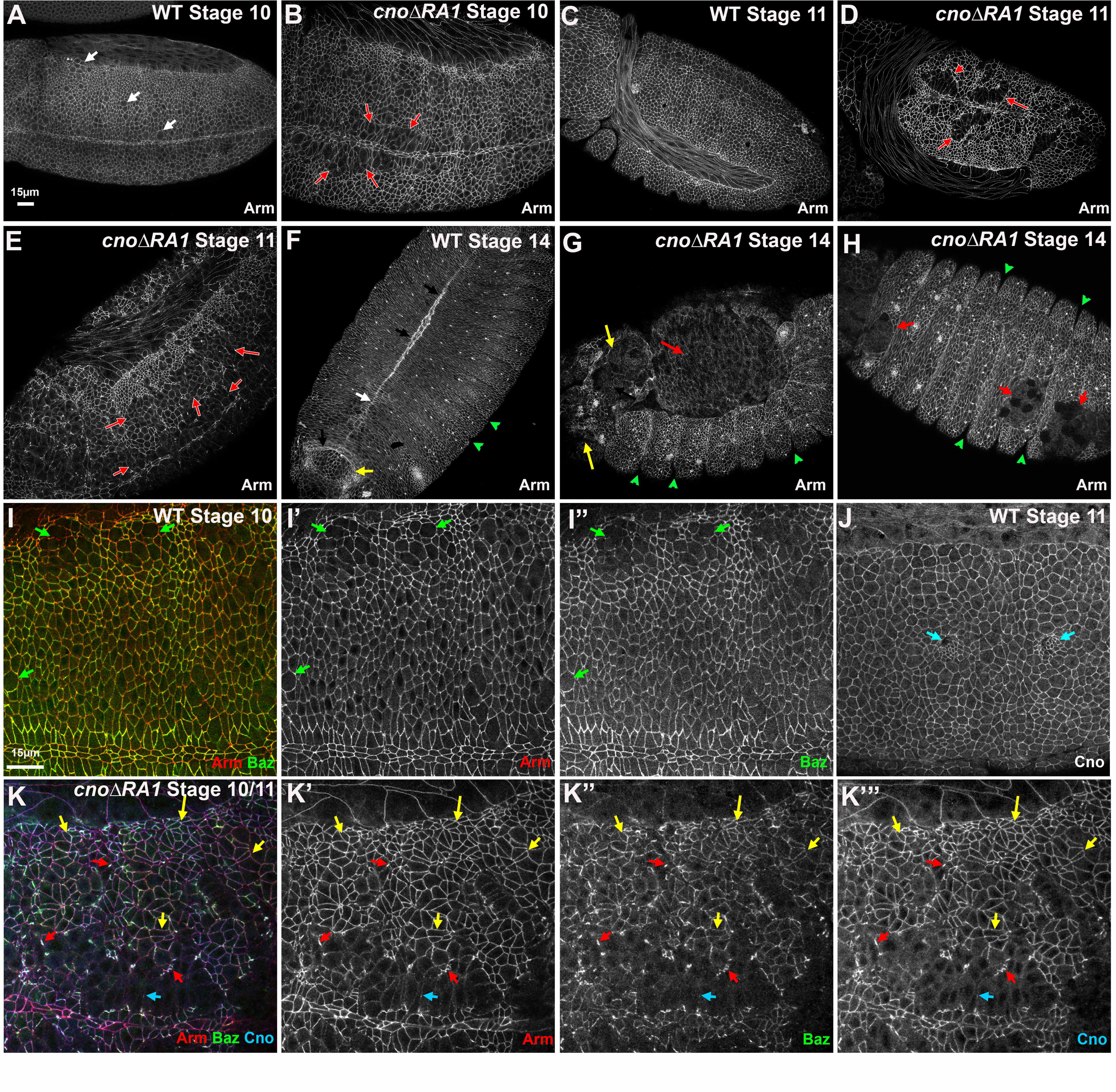
Cno’s RA1 domain is important for dorsal closure, head involution and epidermal integrity. Embryos, anterior left and dorsal up unless otherwise stated, stage, genotype and antigens indicated. (A,B) Stage 10. (A) In wildtype, some cells are rounded up to divide (white arrows), but they rapidly return to columnar architecture. (B) In *cnoΔRA1* mutants, groups of cells near the ventral midline fail to return to columnar architecture (red arrows). (C-E) Stage 11. (C) Wildtype. (D,E) In *cnoΔRA1* mutants, the failure to resume columnar architecture becomes more apparent (red arrows). (F-H) Stage 14 (F) Ventral view. In wildtype dorsal closure is completed (white arrow), head involution is underway (yellow arrow), and segmental grooves are no longer deep (green arrows). (G). *cnoΔRA1* mutant in which dorsal closure has failed, exposing the underlying tissues (red arrow). There are gaps in the head epidermis (yellow arrows) and deep segmental grooves remain (green arrowheads). (H) Ventral view of *cnoΔRA1* mutant with holes in the epidermis (red arrows) and deep segmental grooves (green arrowheads). (I,J) Closeups of wildtype stage 10 (I) and 11(J) showing dividing cells (green arrows) and forming tracheal pits (cyan arrows). Arm, Baz and Cno remain enriched at AJs. (K) Stage 10 *cnoΔRA1* mutant. Cells that have retained columnar architecture retain junctional Arm, Baz and CnoΔRA1 (yellow arrows). However, in some cells junctions appear fragmented (red arrows) and in less epithelial regions Arm, Baz and Cno staining are strongly reduced (cyan arrows).

After germband retraction, two different groups of cells begin complex collective cell migration events. In dorsal closure, the lateral epidermal sheets extend dorsally, and zip together, encasing the embryos in epidermis (Fig. 7F, white arrows), while movements of other more anterior cells complete head involution (Fig. 7F, yellow arrow). In *cnoΔRA1* mutants, both dorsal closure and head involution fail, consistent with the cuticle phenotypes. Dorsal closure is not finished before dorsal amnioserosal cells undergo apoptosis leaving the dorsal side open (Fig. 7G, red arrow), and gaps in epidermal coverage of the head are also apparent (Fig. 7G, yellow arrows). Ventrally, holes remain in the ventral epidermis (Fig. 7H, red arrows), consistent with the earlier defects there (Fig. 7D,E). Persistent deep segmental grooves were seen (Fig. 7G,H green arrows), another *cno* null mutant phenotype. Thus, the RA1 domain is essential for Cno’s roles in dorsal closure, head involution and the maintenance of ventral epidermal integrity.

### A sensitized assay revealed that CnoΔRA2 does not provide full Cno function

The viability and fertility of CnoΔRA2 was somewhat surprising, given its conservation over the ∼600 million years of animal evolution. A subset of our other *cno* mutants, including *cnoΔPDZ* and *cnoΔFAB,* are also viable, despite similar conservation of those protein domains or regions. However, when we used a sensitized assay, we found that neither mutant retains full wildtype function. In this assay, we reduced the maternal and zygotic dose of the mutant protein by making mothers and fathers trans-heterozygous for the mutant of interest and our *cno* null allele, *cno^R2^*. We then examined progeny of this cross. Our control was a cross of parents heterozygous for a wildtype chromosome and *cno^R2^* (*cno^R2^*/+). *cno^R2^* is zygotically embryonic lethal, and thus we expect 25% embryonic lethality in this cross. In parallel we crossed *cno^R2^*/*cnoΔRA2* parents. This led to a modest increase in lethality (35% vs. 28% in the control; Fig. 8A), suggesting that some *cno^R2^*/*cnoΔRA2* embryos were dying.

**Fig. 8.**
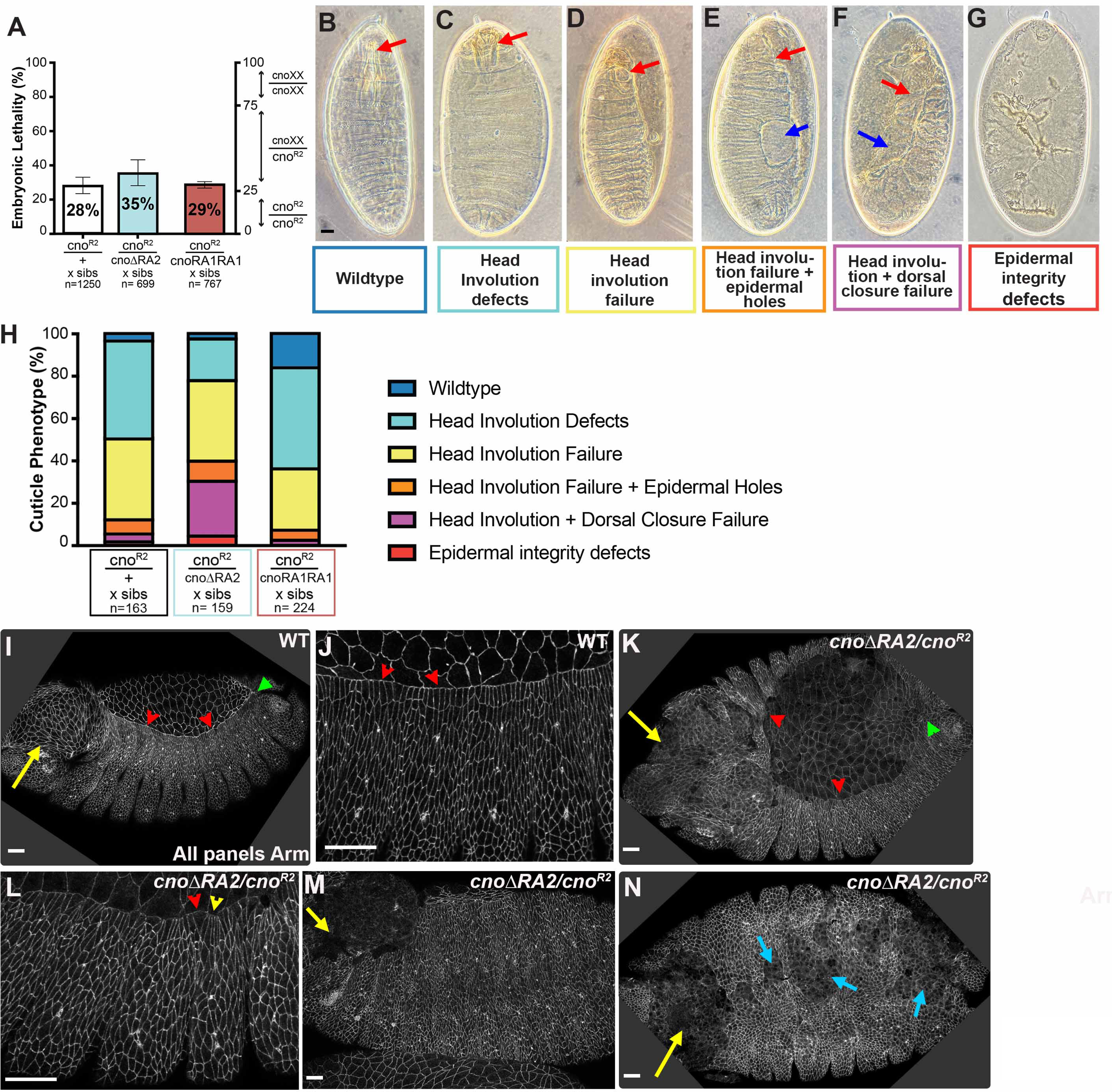
A sensitized assay reveals CnoΔRA2 does not provide fully wildtype function in embryonic morphogenesis. (A) Embryonic lethality. (B-G) Embryonic cuticles illustrating different severity (images are also those used in Fig. 4). (H) Stacked bar graph illustrating the range of defects in different genotypes. Most *cno^R2^* zygotic mutants only have defects in head involution. Progeny of *cno^R2^/cnoΔRA2* parents have more severe defects, with frequent failure of dorsal closure. In contrast, progeny of *cno^R2^/cnoRA1RA1* parents have defects indistinguishable from those of *cno^R2^/+* parents. (I-N) Embryos, anterior left, stages, and antigens indicated. Embryos labeled *cnoΔRA2/cno^R2^* are progeny of the cross of *cnoΔRA2/cno^R2^* parents—genotype was not determined. (I,J) Wildtype stage 13. (I) Dorsal closure (red arrowheads) and head involution (yellow arrow) are proceeding, with zipping beginning at the canthi (green arrow). (J) The leading edge is straight and cell shapes are relatively uniform (red arrowheads). (K-N) *cnoΔRA2/cno^R2^* stage 13-14. The leading edge is no longer straight (K, red arrowheads), zipping is slow at the canthi (K, green arrowhead), leading edge cell shape are less uniform (L, arrowheads), head involution has failed (M,N, yellow arrows), and ventral epidermal holes are sometimes present (N, cyan arrows). Thus, consistent with the cuticle analysis, phenotypes appear stronger. Scale bars =15µm.

We next examined morphogenesis, again using cuticle phenotypes (Fig. 8B-G). Because of the strong maternal contribution of Cno, most events of morphogenesis go to completion in *cno^R2^* zygotic mutants, and most embryos only exhibit defects in head involution (89%; Fig. 8C, D; quantified in 8H), the last major event of epidermal morphogenesis, while dorsal closure generally goes to completion (96% of embryos complete closure). In contrast, the cuticle phenotypes of the embryos from the cross of *cno^R2^*/*cnoΔRA2* parents were substantially enhanced, with 31% of dead embryos exhibiting failure of dorsal closure (n=159). In contrast, *cnoRA1RA1* behaved essentially like a wildtype allele in this assay, with 29% embryonic lethality (Fig. 8A) and morphogenesis defects similar to the *cno^R2^*/+ control: 92% only had defects in head involution and only 3% had defects in dorsal closure (Fig. 8H; n=224).

Consistent with the cuticle phenotypes, we observed defects in dorsal closure and head involution in progeny of the cross of *cno^R2^*/*cnoΔRA2* parents (Fig. 8I-N). We observed defects in zippering and a wavy leading edge during dorsal closure (Fig. 8I vs K, red arrowheads), uneven cell shapes at the leading edge (Fig. 8J vs L) with some cells hyperconstricted (Fig. 8L, yellow arrowhead) and others splayed open (Fig. 8L, red arrowhead). Holes were seen in the head epidermis (Fig. 8I vs K, M, N yellow arrows). These data reveal that CnoΔRA2 does not provide fully wildtype function, in contrast to CnoRA1RA1, which provides full function in this assay.

### Cno’s RA2 domain is required for Cno’s role in patterning the developing eye

In addition to its roles in embryonic morphogenesis, Cno also is important for the intricate cell shape changes and rearrangements that transform the uniform epithelial sheet of the eye imaginal disc into the stereotyped arrangement of epithelial cell types that form each of the ∼750 ommatidia of the mature eye. One way *cno* was first identified was via the effect of weak alleles on the stereotyped arrangement of ommatidial cells (Gaengel and Mlodzik, 2003; Matsuo et al., 1997; Matsuo et al., 1999), while complete loss of function dramatically disrupts epithelial architecture (Walther et al., 2018).

To further explore the role of the RA2 domain, we examined whether *cnoΔRA2* mutants have defects in eye development, examining them 40 h after pupal development has begun. Each nascent ommatidium has a stereotyped arrangement of epithelial cell types, each with characteristic shapes (Fig. 9A,B; (Johnson, 2021). At the center of each ommatidium are four cone cells, surrounded by two primary (1°) pigment cells. These are separated from the cells of neighboring ommatidia by a lattice of rectangular secondary (2°) and hexagonal tertiary (3°) pigment cells, and mechanosensory bristles. Deleting the RA2 domain introduced numerous patterning errors, most notably in the organization of cone cells (Fig. 9C). During eye development cone cells are recruited to ommatidia with the anterior and posterior cells initially organized in direct contact and the dorsal and ventral pair occluded. A T1-T2-T3 junction exchange then reorients this organization, but the anterior-posterior cone cells remained in contact in many *cnoΔRA2* ommatidia, indicating disruptions to this process. We also observed clusters where one of the four cone cells failed to maintain contact with its neighboring 1° cell (Fig. 9C). These defects, as well as minor defects observed in the organization of the lattice cells in *cnoΔRA2* ommatidia were quantified via the ommatidial mis-patterning score (Fig. 9D, Table S1; (Johnson and Cagan, 2009). Molecular control of the T1-T2-T3 transition of cone cells is not well understood, with limited studies suggesting that adhesion rather than changes to the cytoskeleton principally drive the change (Blackie et al., 2021; Grillo-Hill and Wolff, 2009), and our data highlight the importance of Cno in this process. The cone cell defects in *cnoΔRA2* retinas were amplified in *cno^R2^*/ *cnoΔRA2* trans-heterozygotes, where cone cell numbers were occasionally reduced and the lattice was also more severely mis-patterned (Fig. 9E; *cno^R2^*/+ retinas were not mis-patterned, Fig. 9F). We also assessed whether replacing the missing RA2 domain with a second RA1 domain rescued these defects. Surprisingly, cone cell and lattice cell defects were prevalent in *cnoRA1RA1* homozygotes (Fig. 9G), emphasizing an important requirement for the RA2 domain of Cno in pupal eye morphogenesis.

**Fig 9.**
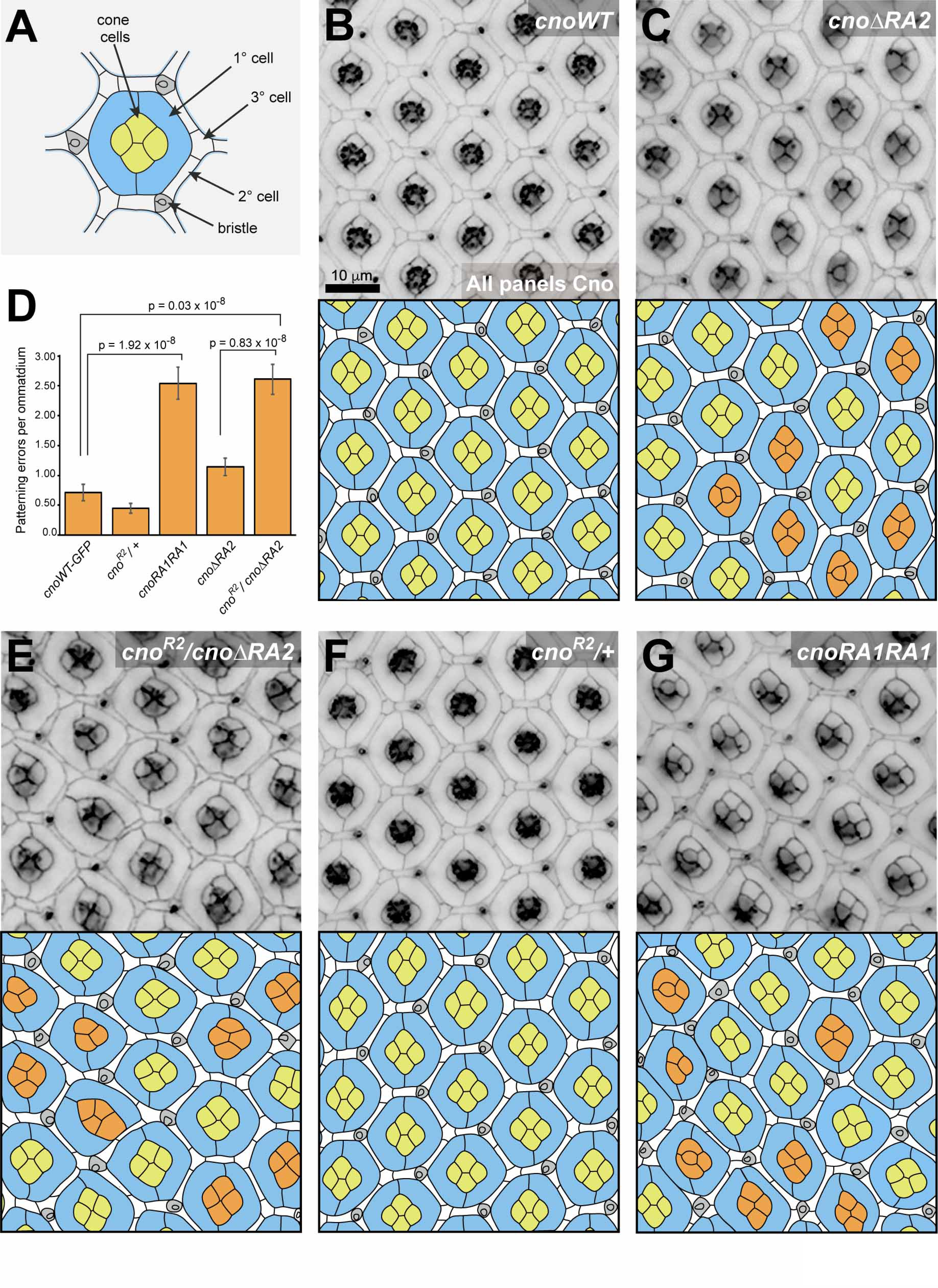
The Cno RA2 domain is essential for cellular morphogenesis in the pupal eye. (A) Cartoon of a pupal eye ommatidium at 40 h APF. (B) Small region of a *cno-wt-GFP* eye and (C) a cno*ΔRA2* homozygous eye, with tracings of the images below. Incorrectly organized cone cells are colored darker green. (D) Patterning errors for all genotypes are quantified as average number of patterning errors per ommatidium, and statistical significance determined using Students T-tests, with p-values as indicated. (E) Small region of *cnoΔRA2/cno^R2^*, (F) *cno^R2^*/+ and (G) *cnoRA1RA1* retinas. GFP-tagged Cno proteins were detected in B, D, E and G, and endogenous Cno in F. Scale bar in B = 10 µm.

## Discussion

Drosophila Cno and mammalian Afadin are key elements of the protein network linking cell-cell AJs to the actomyosin cytoskeleton to allow cells to change shape and move without tissue disruption. Genetic and biochemical data suggest they act as Rap1 effectors in these roles, with Rap1 binding thought to activate Cno and Afadin. However, unlike many effectors of Ras-family GTPases, Cno and Afadin have two RA domains. Deleting both nearly eliminates Cno function, but major questions remain about the biochemical and cell biological functions of the individual RA domains. Here we combine biochemical, genetic, and cell biological analyses to address this important issue.

### RA1 and RA2 have dramatically different roles in Cno’s function in morphogenesis

Strong genetic evidence supports the idea that Cno and Afadin are among Rap1’s effectors, and that Rap1 activates them to carry out their roles in AJ:cytoskeletal linkage (Boettner et al., 2003; Bonello et al., 2018; Hiremath et al., 2023; Pannekoek et al., 2020; Perez-Vale et al., 2023; Sawyer et al., 2009; Walther et al., 2018). Our previous functional assays revealed that deleting both RA domains severely diminished Cno function in multiple events of embryonic morphogenesis (Perez-Vale et al., 2021). However, this left open the question of why Cno and Afadin each have two RA domains. Are they redundant in function or do they have different roles? Our current set of mutants provide a clear answer. RA1 is essential for all of Cno’s functions in embryonic morphogenesis and plays an important role in its recruitment to cell junctions under tension. In contrast, RA2 is dispensable for viability and fertility. It is not completely without a biological function: it is important for high fidelity assembly of the cells that make up each ommatidium in the compound eye, and sensitized assays in embryod reveal it does not provide fully wildtype function. These differential functions contrast with the roles of the two RA domains in PLCε. In this protein, both are essential, with RA2 mediating both PLCε activation by Ras, Rap1 and their upstream activators and its recruitment to the plasma membrane (Bunney et al., 2006; Kelley et al., 2001; Song et al., 2002), and RA1 playing a stabilizing structural role (Rugema et al., 2020). Intriguingly, neither of the RA domains in Cno is essential for recruitment to AJs after gastrulation onset, though both contribute to mechanosensitive recruitment to AJs under elevated tension.

### RA1 and RA2 bind Rap1 with similar affinities

Previous biochemical assays gave different views of the ability of each of the RA domains to bind Rap1. Multiple groups assessed the binding of activated Rap1 to Afadin RA1, measuring Kd’s of 0.2-3µM (Linnemann et al., 1999; Smith et al., 2017; Wohlgemuth et al., 2005). Less quantitative yeast two-hybrid assays also supported binding of Cno RA1 to Rap1 (Boettner et al., 2003). In contrast, the existing data on Afadin or Cno RA2 were conflicting—biochemical assays found no measurable binding between Afadin RA2 and activated Rap1(Wohlgemuth et al., 2005), while yeast two-hybrid assays suggested weaker but detectable binding of Cno RA2 and Rap1 (Boettner et al., 2003). This issue was complicated by the recent realization that RA1 (and likely RA2) has an additional N-terminal alpha-helix that boosts Afadin RA1 binding to Rap1 by 4-fold (Smith et al., 2017). This helix that was missing in the Afadin RA2 construct used to measure binding affinity, and partially truncated in the Cno RA2 constructs used in the two hybrid assays (Boettner et al., 2003).

We thus examined the binding of Cno RA1 or Cno RA2 to activated Rap1, using constructs that retained the N-terminal helix. To our surprise, both SEC-MALs and ITC revealed that each RA domain binds stably to activated Rap1, with similar affinities. This suggests that the dramatic functional differences we observed after deleting each RA domain in vivo are not simply due to only one being able to bind Rap1, or to substantial differences in Rap1 binding affinity. Given the similar affinities, it is truly surprising that they are not at least somewhat redundant in function, but our functional data reveal that is the case. We are thus left having to consider other possibilities. The substantially stronger evolutionary conservation of RA1 between mammals and Drosophila (74% versus 54% amino acid identity for RA2) may suggest that RA1 has an additional role. Perhaps RA1 uses a conserved protein interface outside the Rap1 binding region to interact with other domains of Cno/Afadin or with other protein partners. In the future, identifying what might bind RA1 and not RA2 could provide insights into this issue.

### By what mechanism does Rap1 “activate” Cno?

We have known for more than two decades that Cno acts as a Rap1 effector during embryonic morphogenesis (Boettner et al., 2003). Cno and Rap1 share virtually identical roles in initial positioning of AJs and in mesoderm apical constriction (Choi et al., 2013; Sawyer et al., 2009; Spahn et al., 2012). Rap1 appears to have additional effectors during later embryonic morphogenesis, but data supports the idea that Cno continues to be one of its effectors in germband extension, dorsal closure and epidermal integrity (Boettner et al., 2003; Perez-Vale et al., 2023), as well as during post-embryonic eye development (Walther et al., 2018). We now have confirmed that both Cno RA1 and RA2 bind Rap1 with high affinity, like other effectors of small GTPases. But how does this binding “activate” Cno?

We initially considered two potential roles: Rap1 binding is essential for plasma membrane recruitment and assembly into AJs, or Rap1 binding leads to a conformational change that “opens up” a closed conformation of Cno, allowing it to interact with its junctional and cytoskeletal partners. We tested some of these predictions. The combined RA domains are important for initial polarized recruitment of Cno to nascent AJs as they form during cellularization (Perez-Vale et al., 2021), and Rap1 is also essential for this cortical recruitment. However, CnoΔRA returns to apical AJs as gastrulation starts, and wildtype Cno returns to apical AJs at this time in *Rap1* mutants. Thus, AJ recruitment can occur by multiple means, and this is not Rap1’s sole role. Our earlier data also suggest Rap1 influences Cno localization and function in ways that do not solely rely on the RA domains. Cno is entirely absent from the plasma membrane during junctional assembly in *Rap1* mutants (Sawyer et al., 2009), while CnoΔRA protein remains associated with the lateral membrane but loses its apical polarization (Perez-Vale et al., 2021). This suggests an RA domain-independent effect of Rap1 on Cno’s initial membrane recruitment. Even more puzzling, our new data reveal that the initial recruitment of CnoΔRA1 to the plasma membrane is reduced relative to Cno lacking both RA domains, suggesting that at this stage RA2 may antagonize Rap1-independent membrane recruitment. The multivalent nature of Cno recruitment to AJs is further illustrated by the fact that none of the mutants we have thus far tested, deleting both RA domains, the Dilute domain, the PDZ domain, or the FAB region, abolish gastrulation-stage AJ recruitment of Cno. Recruitment to AJs after gastrulation onset is also Rap1 independent. Thus, AJ recruitment is remarkably robust, and cannot be Rap1’s only role.

In some proteins activated by small GTPase binding, like the formins, GTPase binding triggers a conformational change that releases intramolecular inhibition. This was originally one of our favorite hypotheses. However, in some version of this model, CnoΔRA would be constitutively active, something we did not observe. Further, this idea suggested that the N-terminal folded domains would have a tertiary structure that would be altered by Rap1 binding. The fact that the RA2, Dilute, and PDZ domains are all dispensable for viability has seriously called into question this model. Some versions of this model remain plausible, as RA1 might bind other regions of the protein we have yet to mutate, such as the FHA domain or the IDR. Future mutational analysis, including making versions of RA1 that cannot bind Rap1 but which retain other putative protein interaction interfaces, may provide insights.

## Methods

### Structure prediction and structure analysis

The Drosophila Cno RA domain – Rap1 complex structure prediction models were computed using ColabFold v1.5.5 running AlphaFold2-multimer (Jumper et al., 2021; Mirdita et al., 2022). Sequence inputs were [Canoe RA1 residues 1-137:Rap1 residues 1-184] and [Canoe RA2 residues 222-350:Rap1 residues 1-184]. The mouse Afadin RA1 – human Ras-GMPPNP complex was obtained from PDB 6AMB (Smith et al., 2017). Structures were aligned using the align command in PyMOL (Schrodinger). Protein sequence alignments were generated manually based on output from the PyMOL structural alignments.

### Cloning and purification of Cno RA domains and Rap1 G12V GTPase

DNA encoding the Drosophila melanogaster Cno RA1 and RA2 domains (amino acids 1-137 and 222-350 respectfully) were generated using PCR (RA1 forward primer 5’-CGCTGTTCGCAT ATGTCACATGATAAGAAGATGTTGGATCGCGAGGCAGTACGC-3’; RA1 reverse primer 5’-CACTCGTTCGAATTCTTAGGGAGTGGTCTTTTGGTCTATGTTCTTGAGCAG-3’; RA2 forward primer 5’-GCGTGTTCGCATATGACCCGATCGATCTCAAACCCGGAGGCCGTG-3’; RA2 reverse primer 5’-GAGTCGTTCGAATTCTTATGAGTCCGCCGGACGTCTCCGCA CATG-3’) and sub-cloned into pET28 (MilliporeSigma) using Nde1 and EcoR1 restriction enzymes and T4 DNA ligase (New England Biolabs). Similarly, DNA encoding the Drosophila melanogaster GTPase Rap1 G12V (amino acids 1-180) was generated using PCR and sub-cloned into pET28 using Nde1 and EcoR1 restriction enzymes and T4 DNA ligase (Rap1 forward primer 5’-CGCTGTTCGCATATGCGTGAGTACAAAATCGTGGTCCTTGG-3’; Rap1 reverse primer 5’-CACTCGTTCGAATTCTCATAGGGACTTTTTCGGCTTCTTCTG-3’). The plasmids were transformed into E. coli BL21 DE3 pLysS cells and grown to an optical density at 600 nm of 0.8 in media containing 50 µg/L kanamycin at 37°C. The temperature was lowered to 20 °C and protein expression was induced with 100 µM IPTG for 16 h. Cells were harvested by centrifugation, resuspended in buffer A (25 mM Tris, pH 8.5, 300 mM NaCl, 10 mM imidazole, and 0.1% β-mercaptoethanol [β-ME]) at 4 °C, and lysed by sonication (the Rap1 G12V buffer A was supplemented with 1 mM MgCl2 and 5% glycerol). Phenylmethylsulfonyl fluoride was added to 1 mM final concentration. Cells debris was pelleted by centrifugation at 17,000 × g for one hour and the supernatant loaded onto a 10-ml Ni2+-NTA column (Qiagen). The column was washed with 1 L buffer A and the protein batch eluted with 100 ml buffer B (buffer B = buffer A supplemented with 290 mM imidazole). CaCl2 was added to 1 mM final concentration. RA1 domain protein received 15 µl (3.6 mg/ml) of bovine α-thrombin (Prolytix) and was incubated at 4°C for 36-h to proteolytically cleave off the N-terminal His6 tag. Rap1 G12V protein received 40 µL (9.6 mg/ml) of bovine α-thrombin to proteolytically cleave off the N-terminal His6 tag and was incubated at 4°C for 4 days. Thrombin-treated RA1 and Rap1 G12V batches were dialyzed against 4L of Buffer A with no imidazole for 20 h using 3.5k MWCO SnakeSkin dialysis tubing (ThemoFisher Scientific). After the dialysis step, protein was filtered consecutively over 0.5 ml benzamadine sepharose (Cytiva) and 10-ml Ni2+-NTA column, and the flow through collected. Preparations of RA2, as well as some RA1 preparation, were purified without thrombin treatment or subsequent filtration over benzamidine sepharose or Ni2+-NTA resin. All proteins were dialyzed against 2L of buffer C (25 mM HEPES-NaOH pH 7.5, 100 mM NaCl, 5 mM MgCl2, 0.1% β-ME) for 5 h, followed by a second dialysis against 2L buffer C overnight at 4 °C. Protein was concentrated in a MilliporeSigma molecular weight cutoff centrifugal concentrator (3k MWCO for the RA1 and RA2 constructs; 10k MWCO for the Rap1 G12V construct), aliquoted, and stored at −80°C.

### Nucleotide exchange to generate Rap1-G12V-GMPPNP

30 U of alkaline phosphatase-coupled to agarose beads (MilliporeSigma, P-0762) were washed with 20 mM Tris-HCl pH 7.5. Reaction buffer (40 mM Tris-HCl pH 7.5, 200 mM (NH4)2SO4, 10 µM ZnCl2, 5 mM DTT) was added to beads to bring the volume to 100 µl. To the alkaline phosphatase-coupled agarose bead slurry, 10 mg of Rap1 G12V and 10 M excess GMPPNP (Jena Bioscience) were added and the total volume brought to 1 ml using reaction buffer. The mixture was incubated at 4 °C with rotation for 3 h. The reaction was centrifuged at 2350 x g for 2 min, the supernatant was divided in half, and each half loaded onto a 2-mL Zeba spin desalting column (ThermoFisher Scientific) preequilibrated with buffer C. Spin columns were centrifuged at 1000 x g for 2 min to perform buffer exchange. Protein concentration in buffer C was determined, aliquoted, and stored at -80 °C.

### Biophysical protein analysis

SEC-MALS: Size exclusion chromatography with multiangle light scattering (SEC-MALS) runs were performed in buffer C supplemented with 0.2 g/L of sodium azide. Samples (100 µl) of individual proteins (200 µM) or RA domain – Rap1 G12V GMPPNP complexes (200 µM complex) were prepared in buffer C and injected onto a Superdex 200 column (Cytiva), run at 0.5 ml/min. The eluate was passed over a DAWN light-scattering instrument (Waters | Wyatt Technology Corp.) followed by an Optilab dRI detector (Waters | Wyatt Technology Corp.). Data was processed using the Astra software program (Wyatt Technology Corp.) and plotted using DataGraph (Visual Data Tools).

ITC: Cno RA domain interactions with GMPPNP-loaded Rap1 G12V were measured using an isothermal titration calorimetry (ITC) MicroCal PEAQ-ITC microcalorimeter (Malvern Panalytical). Protein stock solutions of the respective RA domain and GMPPNP-loaded Rap1 G12V were loaded into respective 3.5k MWCO Slide-A-Lyzers (ThermoFisher Scientific) and dialyzed in a shared 1L reservoir of buffer C for one hour, followed by a second 1L reservoir of buffer C for two hours, followed by a third 2L reservoir of buffer C overnight, all at 4 °C. Experiments were carried out at 25 °C. 2-µl volumes of 150-670 µM GMPPNP-loaded Rap1 G12V was automatically injected into a well containing 370 µl of 20-50 µM RA1. 2-µl volumes of 650-1200 µM GMPPNP-loaded Rap1 G12V was automatically injected into a well containing 370 µl of 42-80 µM RA2. 20 injections were performed over the span of one hour with a reference power of 12 µcal/s, an initial delay of 120s, and a stir speed of 750 rpm. Heats of dilution were determined from control experiments in which GMPPNP-loaded Rap1 G12V was titrated into buffer alone. Binding isotherms were fit to a one-site binding model using MicroCal PEAQ-ITC software (Malvern Panalytical).

### Fly work

Unless otherwise noted, all experiments were performed at 25°C. Flies used as controls, referred to in the text as wildtype (WT), were of the *yellow white* genotype (Bloomington Stock #1495). All Cno constructs are listed in Figure 3A. *cnoΔRA1* germline clones were generated by heat shocking 3rd instar larvae generated by crossing *cnoΔRA1/TM3* virgin females to *P{ry[+t7.2]=hsFLP}1, y[1] w[1118]; P{neoFRT}82B P{ovo−D1−18}3R/TM3, ry[*], Sb[1]* males in a 37°C water bath for two hours each on two consecutive days. Then, eclosed virgin females with *hsFLP1; P{neoFRT}82B P{ovo−D1−18}3R/P {neoFRT}82B cnoΔRA1* were collected and subsequently crossed with *cnoΔRA1 / TM3 Sb* males. The embryos generated from this cross were analyzed. This same process was performed to create *cnoRA2RA2* germline clones. Allele viability was assessed via a cross to generate mutant flies that are heterozygous for each *cno* allele with the null allele *cno^R2^* (Sawyer et al., 2009).

### Generating mutant *cno* rescue constructs and their ΦC31-mediated integration into the *cno*ΔΔ allele attP site

The *cnoWT-GFP* rescue construct (Perez-Vale et al., 2021) was used to generate new constructs lacking RA1 (*cnoΔRA1-GFP)*, lacking RA2 (*cnoΔRA2-GFP)*, replacing RA2 with RA1 (*cnoRA1-RA1-GFP)*, and replacing RA1 with RA2 (*cnoRA2-RA2-GFP)*. To generate the *cnoΔRA1-GFP* and *cnoΔRA2-GFP* constructs, amino acids 18-136 (RA1) or 211-353 (RA2) of *Drosophila* Cno were respectively replaced with an ASGGTS polypeptide linker (DNA sequence 5’-GCTAGCGGCGGCACTAGT-3’, which encodes unique NheI and SpeI restriction enzyme sites)—this was done by Azenta Life Sciences (Waltham, MA). To generate the *cnoRA1-RA1-GFP* construct, DNA encoding RA1 (amino acids 1-135) was amplified using PCR, and cloned into the NheI and SpeI sites of the *cnoΔRA2-GFP construct* (forward primer: 5’-CGCATCTGAGCTAGCATGTCACATGATAAGAAGATGTTGGATCGCGAGGCAGTACGC-3’, reverse primer 5’-GCCTAGACTACTAGTGGTCTTTTGGTCTATGTTCTTGAGCAGGAATCGACCTTCGC-3’). To generate the *cnoRA2-RA2-GFP* construct, DNA encoding RA2 (amino acids 211-353) was amplified using the PCR method, and cloned into the NheI and SpeI sites of the *cnoΔRA1-GFP construct* (forward primer: 5’-CGCATCTGAGCTAGCAAACTGTACACGGAACTACCAGAAACCTCGTTCACCCGATCG-3’, reverse primer 5’-GCCTAGACTACTAGTCCGGGGCTGTGAGTCCGCCGGACGTC-3’). All rescue constructs were sequence verified. Each vector carrying the modified *cno* gene, pGE-attB-GMR, also carries a *w^+^* selectable marker next to the *cno* coding sequence, and both are flanked by attR and attL sites allowing site-specific integration into the attP site at the *cnoΔΔ* locus (Perez-Vale et al., 2021). Injection of each mutant *cno*-GFP rescue construct was carried out by BestGene (Chino Hills, CA)— DNA was injected into *PhiC31/int^DM. Vas^; cnoΔΔ* embryos. F1 offspring were screened for the presence of the *w^+^* marker and outcrossed to *w; TM6B, Tb/TM3, Sb* to generate a balanced stock over TM3. We verified the integration of each mutant *cno-GFP* construct by both PCR amplification and sequencing and by immunoblotting. To remove potential other mutations from the mutant *cno-GFP* chromosomes we outcrossed each stock to a *y w* stock with a wildtype 3rd chromosome for multiple generations, selecting for the linked *w^+^* marker in each generation. This allowed us to homozygose *cnoΔRA2-GFP* and *cnoRA1-RA1-GFP.* All mutant *cno-GFP* stocks will be made available via the Bloomington Drosophila Stock Center upon publication.

### Embryo Fixation and Immunofluorescence

Embryo fixation and immunofluorescence were performed as described in McParland *et al*., 2024. Briefly, flies were placed into cups at 25°C with apple juice agar plates containing yeast. Using a paint brush, embryos were collected into 0.1% Triton X-100. Following a five-minute dechorionation with 50% bleach, embryos went through three 15-minute washes in Triton salt solution (0.03% Triton X-100, 68 mM NaCl, EGTA), were fixed at 95°C for 10 seconds, and immediately cooled on ice for at least 30 minutes. Fixed embryos were devitellinized by vigorous shaking in a 1:1 solution of n-heptane and 95% methanol/5% EGTA. Devitellinized embryos were then transferred from the lower methanol layer into new tubes and washed three times with 95% methanol/5% EGTA. Prior to immunostaining, embryos went through three 15-minute washes with phosphate-buffered saline (PBS) with 5% normal goat serum/0.1% saponin (PBSS-NGS) and blocked in PBSS-NGS for 1 hour at room temperature. Antibodies were diluted in a 1% bovine serum albumin/0.1% saponin PBS solution. Following primary antibody incubation either overnight at 4°C or 2-3 hours at room temperature, embryos were washed three times for 15 minutes with PBSS-NGS and incubated in secondary antibodies for 2-3 hours at room temperature. Antibodies and dilutions are in Table 1. Finally, embryos were washed three times with PBSS-NGS and mounted onto glass slides using a homemade Gelvatol solution and stored at 4°C (recipe from the University of Pittsburgh’s Center for Biological Imaging).

**Table 1.** Antibodies Used.

| **Primary Antibodies** | **Host Species** | **Dilution** |
| --- | --- | --- |
| Anti-Armadillo | Mouse IgG<sub>2a</sub> | 1:100 (IF) |
| Anti-α-tubulin | Mouse IgG<sub>1</sub> | 1:5000 (WB) |
| Anti-Bazooka | Rabbit IgG | 1:2000 (IF) |
| Anti-Canoe | Rabbit IgG | 1:1000 (IF)<br>1:2000 (WB) |
| Anti-GFP (JL-8) | Mouse IgG<sub>2a</sub> | 1:1000 (WB) |
| Anti-GFP | Chicken IgY | 1:1000 (IF) |
| **Secondary Antibodies** | **Host Species** | **Dilution** |
| Anti-Mouse IgG<sub>2a</sub> Alexa Fluor 647 | Goat IgG | 1:1000 (IF) |
| Anti-Rabbit IgG Alexa Fluor 568 | Goat IgG | 1:1000 (IF) |
| Anti-Chicken IgY Alexa Fluor 488 | Goat IgG | 1:1000 (IF) |
| Anti-Rabbit IgG IRDye 680RD | Goat IgG | 1:10000 (WB) |
| Anti-Mouse IgG IRDye 800CW | Donkey IgG | 1:10000 (WB) |

### Image Acquisition and Analysis

Following fixation and immunostaining, embryos were imaged on an LSM 880 confocal laser-scanning microscope using a 40x/NA 1.3 Plan-Apochromat oil objective (Carl Zeiss, Jena Germany). ZEN 2009 software was used to acquire and process images and generate maximum intensity projections. Input levels, brightness, and contrast were adjusted using Photoshop (Adobe, San Jose, CA). Apical-basal positioning on maximum intensity projections (MIPs) was analyzed based on Choi *et al,* 2013. Z-stack outputs (at 1.6 digital zoom) were cropped to 200x200 pixels in the region of interest in ZEN 2009 software. Dimensions of the ROIs were converted from *yzx* to *xyz,* and MIPs were generated.

### Cno SAJ and TCJ/multicellular junction enrichment analysis

Images of z-stacks taken through the embryo with a digital zoom of 1.6 or 2 and a step size of 0.3 µm were used for spot adherens junction (SAJ) and TCJ enrichment analysis. Cno TCJ intensity was analyzed in MIPs in stage 7 embryos. MIPs were generated in FIJI (National Institutes of Health, Bethesda, MD, USA) from the apical 1.2-2.4 µm region of z-stacks with a digital zoom of 1.6 or 2 and a step size of 0.3 µm. Lines were drawn (width of 5 pixels) in FIJI to measure mean Cno intensity of bicellular (avoiding multicellular junctions), TCJs/multicellular junctions, and cytoplasm background signal at a 300% digital zoom. Cytoplasmic background was subtracted from all junctional signals to standardize pixel intensity for individual junctions. To calculate the Cno TCJ ratio, the mean standardized TCJ/multicellular junction intensity was divided by the mean standardized bicellular junction intensity (calculated from the average of all bicellular junctions in each TCJ). Ten multicellular junctions were analyzed per embryo yielding box and whisker plots showing 25th-75th percentile ratios in the box and whiskers showing 5th-95th percentiles. Graphs were produced and data statistical analyses were performed in GraphPad Prism version 10.0.0 (GraphPad Software, Boston, Massachusetts USA) using Welch’s unpaired t-test or Brown Forsythe and Welch ANOVA test.

### Planar polarity quantification

Stage 7 to early stage 8 embryo apical 1.2-2.4 µm regions were measured to obtain planar polarity of Cno between anteroposterior (AP) and dorsoventral (DV) borders. Similar to multicellular junction analysis, lines were drawn (width of 5 pixels) at 300% zoom at bicellular borders splitting AP and DV axes and cytoplasm background signal to quantify pixel intensity using FIJI. Cytoplasmic background signal was subtracted from bicellular border intensity to generate standardized Cno intensity. Average AP border standardized Cno intensity was divided by average DV border standardized Cno intensity to generate an AP/DV Cno ratio to quantify planar polarity. Ten AP and ten DV borders were measured per embryo yielding box and whisker plots showing 25th-75th percentile ratios in the box and whiskers showing 5th-95th percentiles. Graphs were produced and data statistical analyses were performed in GraphPad Prism version 10.0.0 (GraphPad Software, Boston, Massachusetts USA) using Welch’s unpaired t-test or Brown Forsythe and Welch ANOVA test.

### Cuticle preparation and analysis

We prepared embryonic cuticles as in Wieschaus and Nüsslein-Volhard (1986). Embryos were collected from apple juice agar plates with yeast and transferred to apple juice agar plates that incubated for at least 48 hours at 25°C. Unhatched embryos were tallied and collected into 0.1% Triton X-100 where they were then dechorionated in 50% bleach for 5 minutes, and washed three times with 0.1% Triton X-100. Unhatched embryos were transferred onto glass slides and mounted in a 1:1 solution of Hoyer’s/lactic acid (recipe from Cold Springs Harbor Protocols) and incubated at 60°C for 24-48 hours and then stored at room temperature. Images were captured on an iPhone using a Nikon Labophot with a 10x Phase 2 lens and categorized based on morphological criteria.

### Immunoblotting

Table 1 contains the antibodies and dilutions used for these experiments. Expression levels of Cno, GFP, and α-tubulin proteins were determined by immunoblotting embryonic lysates that were collected at 1-4 hours and 12-15 hours timepoints. Embryonic lysates were generated as in Manning *et al*., 2019. Briefly, embryos were collected into 0.1% Triton X-100, dechorionated for 5 minutes in 50% bleach, and thrice washed with 0.1% Triton X-100. Subsequently, lysis buffer (1% NP-40, 0.5% Na deoxycholate, 0.1% SDS, 50 mM Tris pH 8.0, 300 mM NaCl, 1.0 mM DTT, Halt^TM^ Protease and Phosphatase Inhibitor Cocktail (Thermo Fisher Scientific, #78442) (100×), and 1 mM EDTA) was added, and embryos were ground with a pestle for ∼20 seconds and placed on ice. After 10 minutes, embryos were again ground with a pestle for ∼20 seconds and subsequently centrifugated at 16,361 *g* for 15 minutes at 4°C. Protein concentration was assessed using Bio-Rad Protein Assay Dye and recording absorbance at 595 nm with a spectrophotometer. After resolving lysates using 7% SDS-PAGE, proteins were transferred onto nitrocellulose membranes. Prior to immunostaining, blocking was performed using 10% bovine serum albumin (BSA) diluted in Tris-buffered saline with 0.1% Tween-20 (TBST) for 1 hour at room temperature. Primary and secondary antibodies were diluted in 5% BSA diluted in TBST. Primary antibody incubation was performed overnight at 4°C, and secondary antibody incubation was performed for 45 minutes at room temperature. The Odyssey CLx infrared system (LI-COR Biosciences) was used to image the membranes, and band densitometric analysis was performed using Empiria Studio® Software (LI-COR Biosciences).

### Pupal eye dissection, immunofluorescence and analysis

Fly cultures were maintained at 25°C on nutrient-rich Drosophila media and pre-pupae selected and maintained in humidified chambers until dissection at 40 hours after puparium formation (h APF) {DeAngelis, 2019 #26}. Rabbit anti-Cno (1:500) or chicken anti-GFP (1:8000, Abcam #13970) were used to detect Cno, with secondary antibodies obtained from Jackson ImmunoResearch. Dissections were performed in triplicate, with 5-10 pupae of each genotype dissected each time. Images were gathered with a Leica DM5500 B fluorescence microscope, processed for publication using Adobe Photoshop and patterning errors were scored in retinas 8-14 retinas for each genotype as previously described (Johnson and Cagan, 2009). Students’ T-tests were used to determine statistical differences between the mis-patterning of different genotypes.

## Supplemental Material

**Fig. S1.**
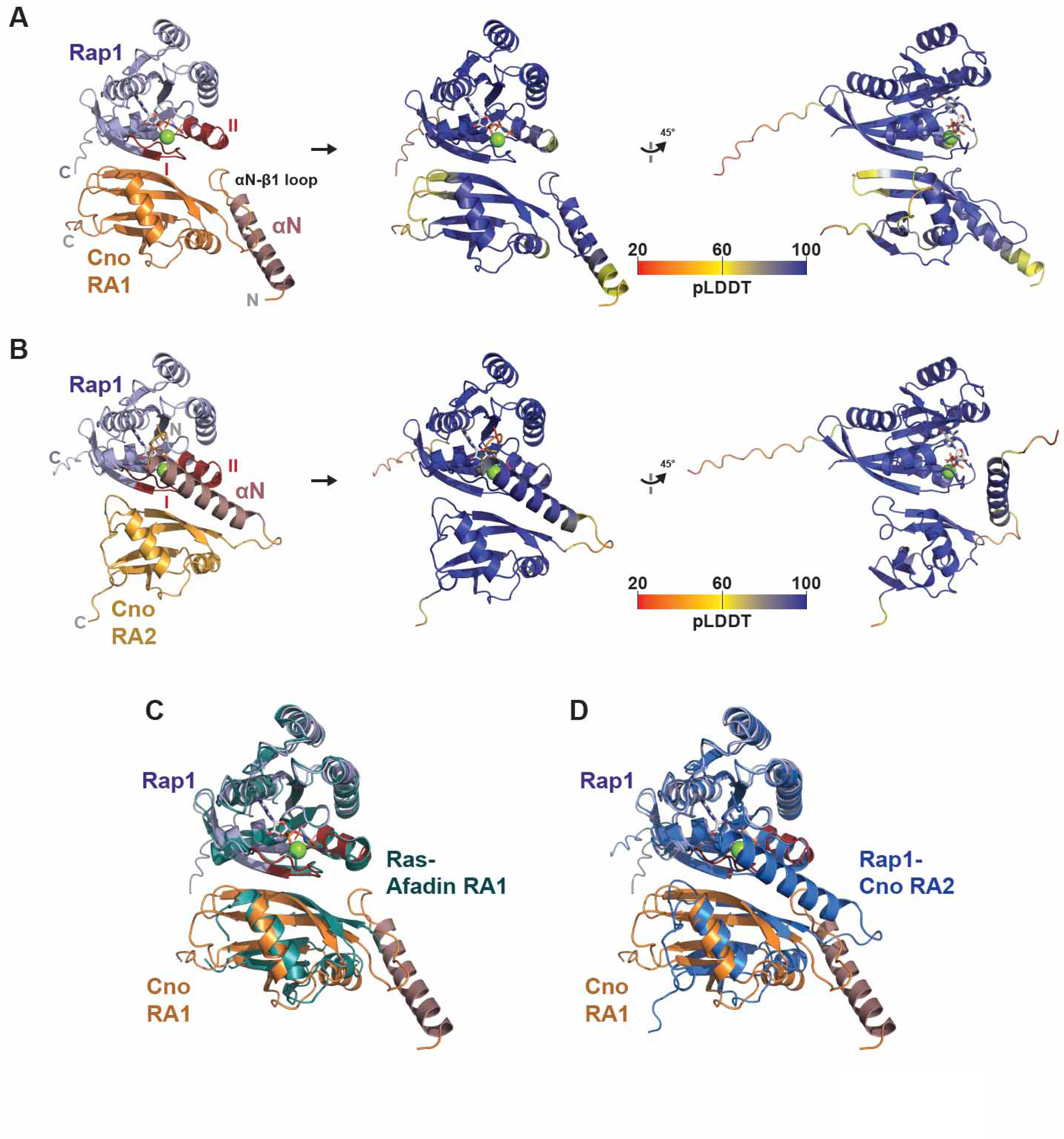
AlphaFold2 structure predictions of Cno RA domain:Rap1 complexes have high confidence and align with the crystal structure of Afadin RA1:Ras. (A,B) AlphaFold2 structure models of the D.m. Cno RA1:Rap1 complex (A) and the D.m. Cno RA2:Rap1 complex (B) shown in cartoon format at left (as shown in Fig. 1 C,D), and colored according to prediction confidence represented by pLDDT value, right two panels, undergoing rotation as shown. (C) Structure alignment of the D.m. Cno RA1:Rap1 AlphaFold2 structure prediction and the mouse Afadin RA1:human Rap1-GMPPNP complex crystal structure (PDB ID 6AMB) (RMSD = 0.70 Å over 147 Cα atoms; (Smith et al., 2017). (D) Structure alignment of the D.m. Cno RA1:Rap1 and RA2:Rap1 AlphaFold2 structure predictions (RMSD = 1.08 Å over 204 Cα atoms).

**Figure S2.**
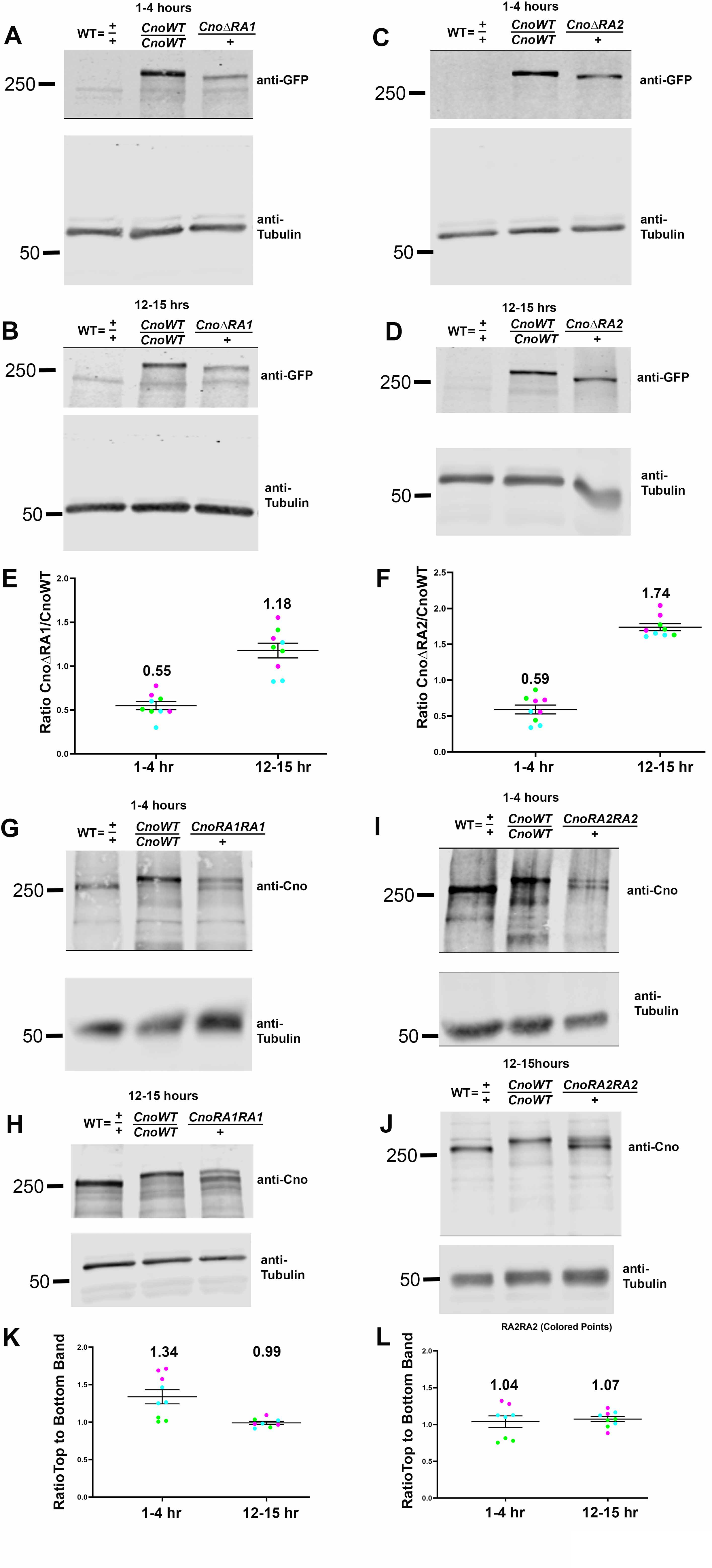
Our Cno mutant proteins accumulate levels similar to wildtype. (A,B,C,D) Embryonic protein extracts of the indicated genotypes and timepoints, immunoblotted with antibodies to GFP, with α-tubulin as an internal loading control. The individual RA domains have a similar molecular weight to the C-terminal GFP fusion, thus causing GFP-tagged CnoΔRA1 and CnoΔRA2 proteins to run at an apparent molecular weight similar to wild-type Cno. Thus, for these genotypes we compared levels using the GFP-antibody, with cnoWT-GFP, which we previously found accumulates at levels similar to wildtype Cno, as our standard for quantifying mutant protein accumulation. (E,F) Calculated levels of CnoΔRA1 and CnoΔRA2 relative to CnoWT. Band intensity values for CnoΔRA1 and CnoΔRA2 were multiplied by a factor of two to account for copy number difference compared to CnoWT. Technical replicates (dots) for each biological replicate (colors) are shown, with the wider band illustrating the mean value and the narrower bands representing the s.e.m. (G,H,I,J) Embryonic protein extracts of the indicated genotypes and timepoints, immunoblotted with antibodies to the C-terminus of Cno and α-tubulin as an internal loading control. *yellow white* (WT) and cnoWT-GFP (CnoWT) embryos were positive controls for Cno antibody. (K,L) Calculated levels of CnoRA1RA1 and CnoRA2RA2 relative to wild-type Cno. Due to sufficient separation between mutant and wild-type Cno bands, quantitation of CnoRA1RA1 and CnoRA2RA2 accumulation was achieved by evaluating the ratio of mutant Cno (top band) to wild-type Cno (bottom band). Technical replicates (dots) for three biological replicates are shown, with the wider band illustrating the mean value and the narrower bands representing the s.e.m.

**Table S1:** Patterning defects per ommatidium (N=76 to 110 ommatidia per genotype)

| Genotype | Cone cell defects |  | Primary cell defects |  | Ommatidial misorientation |  | Bristle defects |  | Tertiary cell defects |  | Number of lattice cells |  | Mis-patterning score |  |  | Mis-patterning score comparison (p-value, * = significant) |
| --- | --- | --- | --- | --- | --- | --- | --- | --- | --- | --- | --- | --- | --- | --- | --- | --- |
|  | Mean | Std Dev | Mean | Std Dev | Mean | Std Dev | Mean | Std Dev | Mean | Std Dev | Mean | Std Dev | Mean | Std Dev | Std Error |  |
| *w<sup>1118</sup>; cno-wt::GFP* | 0.00 | 0.00 | 0.00 | 0.00 | 0.00 | 0.00 | 0.20 | 0.40 | 0.25 | 0.47 | 11.95 | 0.51 | 0.71 | 1.20 | 0.14 |  |
| *w<sup>1118</sup>; cno<sup>R2</sup> / +* | 0.04 | 0.21 | 0.00 | 0.00 | 0.00 | 0.00 | 0.03 | 0.18 | 0.08 | 0.27 | 12.27 | 0.54 | 0.45 | 0.78 | 0.08 | 0.1063 compared to *cno-wt::GFP* |
| *w<sup>1118</sup>; cnoRA1RA1::GFP* | 1.06 | 1.12 | 0.12 | 0.40 | 0.01 | 0.11 | 0.27 | 0.45 | 0.35 | 0.60 | 11.75 | 1.03 | 2.54 | 2.44 | 0.27 | 1.9209 x 10<sup>-8</sup> * compared to *cno-wt::GFP* |
| *w<sup>1118</sup>; cno-ΔRA2::GFP* | 0.25 | 0.54 | 0.10 | 0.56 | 0.00 | 0.00 | 0.13 | 0.34 | 0.22 | 0.42 | 11.69 | 0.76 | 1.14 | 1.43 | 0.14 | 0.0313 compared to *cno-wt::GFP* |
| *w<sup>1118</sup>; cno-ΔRA2::GFP / cno<sup>R2</sup>* | 0.82 | 0.75 | 0.25 | 0.81 | 0.08 | 0.27 | 0.34 | 0.53 | 0.41 | 0.59 | 11.45 | 0.90 | 2.61 | 2.62 | 0.25 | 0.8291 x 10<sup>-8</sup> * compared to *w<sup>1118</sup>; cno-ΔRA2::GFP*<br>0.0265 x 10<sup>-8</sup> * compared to *cno-wt::GFP* |

